# Evolution of barrier loci at an intermediate stage of speciation with gene flow

**DOI:** 10.1101/703108

**Authors:** Xiaodong Liu, Sylvain Glémin, Sophie Karrenberg

## Abstract

Understanding the origin of new species is a central goal in evolutionary biology. Diverging lineages often evolve highly heterogeneous patterns of differentiation; however, the underlying mechanisms are not well understood. We used an integrated approach to investigate evolutionary processes governing genetic differentiation between the hybridizing campions (*Silene dioica* (L.) Clairv. and *S. latifolia* Poiret). Demographic modeling indicated that the two species diverged with continuous gene flow. The best-supported scenario with heterogeneity in both migration rate and effective population size suggested that 5% of the loci evolved without gene flow. Differentiation (*F*_ST_) and sequence divergence (*d*_XY_) were correlated and both tended to peak in the middle of most linkage groups, consistent with reduced gene flow at highly differentiated loci. Highly differentiated loci further exhibited signatures of selection and differentiation was significantly elevated around previously identified QTLs associated with assortative mating. In between-species population pairs, isolation by distance was stronger for genomic regions with low between-species differentiation than for highly differentiated regions that may contain barrier loci. Moreover, differentiation landscapes within and between species were only weakly correlated suggesting that the interplay of background selection and conserved genomic features is not the dominant determinant of genetic differentiation in these lineages. Instead, our results suggest that divergent selection drove the evolution of barrier loci played and the genomic landscape of differentiation between the two species, consistent with predictions for speciation in the face of gene flow.

## Introduction

Understanding the origin of new species is a central goal in evolutionary biology. Genetic differentiation between diverging lineages often is highly heterogeneous throughout the genome; however, the underlying causes are much debated (Ravinet et al., 2017; Wolf & Ellegren, 2017) and empirical studies are still scarce, particularly for intermediate stages of speciation. One important determinant of the evolutionary processes at work is the demographic history of lineage divergence (Ravinet et al., 2017). When geographic barriers prevent gene flow, drift and adaptation can proceed independently in each lineage. Under this scenario, high differentiation may commonly arise in regions of low recombination, through the ubiquitous process of background selection (Burri, 2017; Burri et al., 2015; Nachman & Payseur, 2012). Under a contrasting scenario, speciation occurs with ongoing gene flow. Here, regions of high differentiation may arise because of reduced gene flow at barrier loci whereas the remainder of the genome is homogenized by gene flow. Theoretical studies indicate that divergence with gene flow is most likely when loci controlling reproductive barriers are clustered in the genome or situated in regions of low recombination (Butlin & Smadja, 2018; Yeaman, Aeschbacher, & Bürger, 2016). A third common scenario combines divergence in allopatry with gene flow upon secondary contact. In this case, gene flow erodes differentiation at loci unlinked to adaptive differentiation or reproductive isolation. This situation may also prompt the evolution of further reproductive barriers (reinforcement) when intermediate hybrids are selected against (Butlin & Smadja, 2018). More complex scenarios, including effects of changes in population size and recurrent bouts of gene flow may be also supported (Christe et al., 2017). Divergent selection, background selection and gene flow leave distinct population genomic signatures (see below); however, it is clear that realistic demographic scenarios are a pre-requisite to interpret the genomic landscape of differentiation.

Despite the strong evidence for variation in gene flow throughout the genome and pervasive effects of background selection and conserved genomic features, such as recombination rate variation, mutation rate and gene density, on genetic differentiation (Ravinet et al., 2017; Wolf & Ellegren, 2017), demographic models have only just begun to incorporate these heterogeneities (Christe et al., 2017; Rougemont & Bernatchez, 2018; Roux et al., 2014; Roux et al., 2016; Tine et al., 2014). Models based on diffusion approximation (Gutenkunst, Hernandez, Williamson, & Bustamante, 2009) or on approximate Bayesian computation (Roux et al., 2016) of the joint site frequency spectrum can explicitly include heterogeneous migration rates. Heterogeneities in conserved genomic features and thus background selection, on the other hand, can be modeled by allowing effective population size, *N_e_*, to vary among loci (Charlesworth, 2009; Roux et al., 2016). Including heterogeneity in migration rate and *N_e_* in demographic models often improves model fit and can affect inferences on the best-supported model as well as demographic estimates, such as divergence time (Rougemont & Bernatchez, 2018; Roux et al., 2016). Interestingly, support for heterogeneous migration rates in demographic models was strongest in taxon pairs at intermediate stages of speciation, species, in the so-called “grey zone” of speciation (Roux et al., 2016). This finding aligns with empirical results on the genomic landscape of differentiation, reviewed in Ravinet et al. (2017) and Wolf and Ellegren (2017). At later stages of speciation, gene flow may rapidly cease leading to a genome-wide rise of differentiation (Flaxman, Wacholder, Feder, & Nosil, 2014). The evolution of heterogeneous genetic differentiation is key to understanding the speciation process and it therefore is most promising to target taxa that have considerable reproductive isolation but are still hybridizing.

Population genomic signals of gene flow, divergent selection, and background selection on the genomic landscape of differentiation have been the subject of recent research (Burri, 2017; Cruickshank & Hahn, 2014; Ravinet et al., 2017; Wolf & Ellegren, 2017). Genetic differentiation is commonly measured using Wright’s fixation index, *F*_ST_, which is sensitive not only to changes in gene flow, but also to alterations of genetic diversity within lineages (Burri, 2017; Cruickshank & Hahn, 2014; Wolf & Ellegren, 2017). Sequence divergence, often measured as *d*_XY_, in contrast, is expected to increase primarily due to reductions in gene flow (barrier loci) or as a result of lineage sorting in ancestral populations (Nachman & Payseur, 2012; Richards, Servedio, & Martin, 2019). Joint increases in *d*_XY_ and *F*_ST_ are therefore expected at barrier loci but not at loci with high differentiation due to background selection (Cruickshank & Hahn, 2014; Nachman & Payseur, 2012). Several studies have further inferred an important role of divergent selection in the generation of highly differentiated loci by testing for reductions in sequence diversity and for an excess of rare variants (Nielsen, 2005), for example, in three-spined sticklebacks (Feulner et al., 2015; Marques et al., 2017; Samuk et al., 2017), cichlids (Malinsky et al., 2015; Meier, Marques, Wagner, Excoffier, & Seehausen, 2018) and poplars (Wang, Street, Scofield, & Ingvarsson, 2016). Moreover, admixture analyses and analyses of multiple population pairs in different geographic contexts have identified genomic regions with reduced gene flow, as in sea bass (Duranton et al., 2018) and Darwińs finches (Han et al., 2017). Thus, even though many of these approaches remain challenging, the combined use of different population genomic estimates and analyses helps to elucidate processes affecting genetic differentiation.

Ideally, speciation genomic studies relate to well-investigated reproductive barriers and the genes controlling them. The nature, strength and genetic architecture of reproductive barriers determine the course and completion of the speciation process (Flaxman et al., 2014). Some speciation events appear to be based on few traits controlled be a small number of genomic regions, for example in crows (Poelstra et al., 2014). In many other systems, in contrast, speciation can proceed through multiple reproductive barriers, including adaptive differentiation, assortative mating, and intrinsic barriers, which may be associated with a large number of traits and/or complex genetic architectures (reviewed in Wolf & Ellegren, 2017). In cases involving many loci of small effect, it may be difficult or impossible to identify the genes underlying reproductive isolation. This is because the genetic architecture of reproductive isolation may be transient and the effect of individual loci may be too small to be detected (Rockman, 2012; Yeaman, 2015). Nonetheless, integrating the ecology and genetic control of reproductive barriers with population genomic analyses is necessary to understand the speciation process.

We investigated the closely related campions, *Silene dioica* (L.) Clairv. and *S. latifolia* Poiret (Caryophyllaceae), a plant system with ongoing hybridization but near-complete reproductive isolation (reproductive isolation index, RI_total_ > 0.98) (Karrenberg & Favre, 2008; Karrenberg et al., 2019; Minder, Rothenbuehler, & Widmer, 2007). The two species have separates sexes with XY sex determination and largely overlapping European distributions (Friedrich, 1979). Reproductive isolation mainly results mainly from adaptation to different habitats and from pollinator-mediated assortative mating (Favre, Widmer, & Karrenberg, 2017; Goulson & Jerrim, 1997; Karrenberg et al., 2019). Pink-flowered *S. dioica* occurs in colder habitats, whereas white-flowered *S. latifolia* is found in in warmer and more disturbed habitats (Friedrich, 1979; Karrenberg & Favre, 2008). Intrinsic and post-zygotic barriers, such as pollen competition and hybrid breakdown, are comparatively weak suggesting that ecological divergence drives speciation (Favre et al., 2017; Karrenberg et al., 2019). The genetic architecture of traits associated with various reproductive barriers involves a large number of loci distributed over most of the genome, with evidence for clustering (Liu & Karrenberg, 2018). Hu and Filatov (2015) estimated net non-synonymous sequence divergence (*D*_a_) between the species to 0.0027 for autosomal loci, similar to animal systems with incomplete speciation (Roux et al., 2016). Demographic models further suggest that divergence with gene flow is more likely than strict isolation (Guirao-Rico, Sánchez-Gracia, & Charlesworth, 2017; Hu & Filatov, 2015; Muir, Dixon, Harper, & Filatov, 2012); however, secondary contact scenarios have not been tested and heterogeneity in migration rates or effective population size was not modeled thus far.

In this study, we investigated evolutionary processes driving genetic differentiation between *Silene dioica* and *S. latifolia* and using range-wide population sampling and a reduced representation sequencing technique (double-digest RAD sequencing; Peterson, Weber, Kay, Fisher, & Hoekstra, 2012), together with previous results on QTL mapping for traits associated with reproductive barriers (Liu & Karrenberg, 2018). We first explored the demographic history of the two species taking into account heterogeneity in migration rate and effective population size. As a second step, we described the genomic landscape of differentiation using linkage maps (Liu & Karrenberg, 2018) and tested for signatures of selection in highly differentiated regions. Moreover, we investigated whether regions with elevated differentiation are associated with QTLs for reproductive barrier traits (Liu & Karrenberg, 2018) and with reduced gene flow. This integrated approach allows us to evaluate evidence for barrier loci and infer the evolutionary forces generating them.

## Materials and methods

### Sampling of species and populations

We sampled 11 populations of *Silene dioica* and 9 populations of *S. latifolia* throughout their distribution ranges (Figure 1a). For each population, maternal seed families were collected from individuals at least 5 m apart. Seeds of 8-12 maternal families per population, except for the *S. latifolia* population from Russia (RUS1), which was sampled as a pooled seed lot without family structure, were grown in the greenhouse at Evolutionary Biology Centre, Uppsala University, Sweden (Table S1, Supporting Information). 189 individuals of *S. dioica* and 162 of *S. latifolia* were sampled for DNA extraction and ddRADseq sequencing.

**Figure 1.**
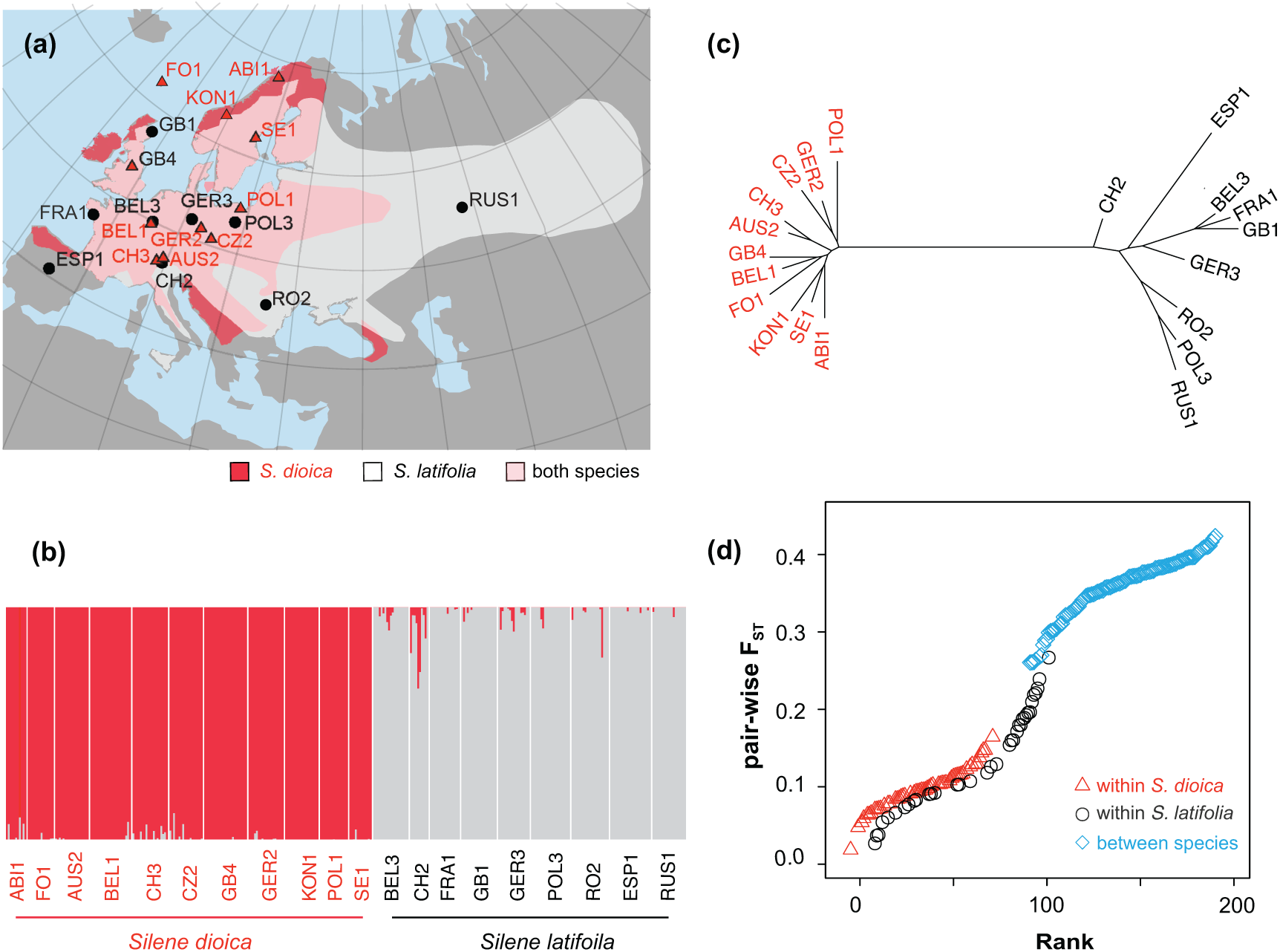
**(a)** Geographic distribution of *Silene dioica* (triangles, 11 populations) and *S. latifolia* (circles, 9 populations); **(b)**, barplot of a structure analysis with *K* = 2, the number of species included in the study; **(c)**, unrooted phylogenetic tree using a neighbor-joining method based on Nei’s genetic distance; **(d),** ranked *F*_ST_ values between population pairs within *S. dioica,* within *S. latifolia* and between the two species.

### ddRAD sequencing and genotyping

Genomic DNA from silica-dried leaf tissues was extracted using Qiagen’s DNeasy Plant Mini Kit (Qiagen, Germany) and quantified using a Qubit dsDNA HS Fluorometer (Life Technologies, Sweden). We prepared libraries for double-digest RAD sequencing (ddRAD-seq) with the restriction enzymes EcoRI and Taq^α^I as described in Liu and Karrenberg (2018). Briefly, enzymatically digested DNA was ligated with barcoded adaptors and size-selected to approximately 550 bp (Peterson, Weber, Kay, Fisher, & Hoekstra, 2012). In total, nine 48-plex libraries were sequenced on Illumina HiSeq 2500 systems at the SNP&SEQ technology platform of SciLifeLab, Uppsala, Sweden using 125-bp paired-end chemistry with two libraries combined to one lane.

ddRAD-seq data were processed following the dDocent pipeline v2.2 (Puritz, Hollenbeck, & Gold, 2014). First, we de-multiplexed raw reads using the *process_radtag* function of Stacks (Catchen, Hohenlohe, Bassham, Amores, & Cresko, 2013). We then pruned bases of low quality and adapter sequences by Trimmomatic (Bolger, Lohse, & Usadel, 2014). We implemented BWA-MEM v0.7.16 (Li, 2013) to align cleaned reads to reference contigs previously assembled from eight deeply sequenced individuals from Switzerland (Liu & Karrenberg, 2018), as there is no full genome sequence available for *Silene* thus far (Krasovec, Chester, Ridout, & Filatov, 2018). We extended the reference with contigs built from unmapped pairs of reads with occurrences of at least 4X within an individual and present in at least 4 individuals using the *de novo* RAD assembler Rainbow (Chong, Ruan, & Wu, 2012). We aligned reads to the new reference with BWA-MEM v0.7.16 allowing up to 3 mismatches and excluded highly clipped sequences (Li, 2013). We employed Freebayes (Garrison & Marth, 2012) for variant calling on the basis of populations with a minimum mapping quality score of 5 and a minimum base quality of 5.

Raw variants were filtered using VCFtools (Danecek et al., 2011) with the following criteria: a minimum quality score of 30, a minimum coverage of 6X, and a minimum genotype call rate of 70% across all samples excluding those were genotyped at < 6% of all sites. We later decomposed complex variant calls into single SNPs using the *vcfallelicprimatives* command from vcflib (Garrison, 2012). To filter spurious SNPs potentially due to paralogs, we excluded SNPs at which allele balance (AB) for heterozygous genotypes was below 0.25 or above 0.75, SNPs covered by both forward and reverse reads and SNPs with excessive coverage (> 100X) (O’Leary, Puritz, Willis, Hollenbeck, & Portnoy, 2018). We included rare variants, which potentially evolved recently, to avoid biased estimation of site frequency spectra (no filter for minor allele frequency).

### Population structure

We analyzed population structure using ADMIXTURE (Alexander, Novembre, & Lange, 2009).To prepare the input, SNPs in high linkage disequilibrium were removed using PLINK (Purcell et al., 2007) with the recommended settings: a window size of 50 SNPs, a step size of 10 SNPs and threshold of *R*^2^ equal to 0.1. We examined the clustering of individuals in ADMIXTURE with the number of groups (*K*), set to two, the number of species, and 20, the number of populations. In addition to the ADMIXTURE analysis, we also assessed the relationships between populations using a neighbor-joining tree based on genetic distance in the R package *hierfstat* (Goudet, 2004).

### Demographic modeling

To investigate the evolutionary history of lineage-split between *Silene. dioica* and *S. latifolia,* we performed demographical modeling based on the folded joint site frequency spectrum (SFS) using the software ∂a∂i, which implements an diffusion approximation-based approach (Gutenkunst et al., 2009). We randomly selected one SNP per reference contig for subsequent analysis in order to satisfy the condition of independence among SNPs for composite likelihood ratio tests. To avoid close relatedness among individuals, we also randomly picked two individuals of different families per population. The SFS was projected onto 20 haploid samples, corresponding to 10 diploid individuals in each group to maximize the number of segregating sites.

We first considered a standard set of demographic scenarios: strict isolation without gene flow (SI), isolation with gene flow (IM), secondary contact (SC) and ancient migration (AM) (Figure S2, Supporting Information). We added population expansion to each model (“exp” models). In separate models, we added heterogeneity in migration rate, *m*, (for models with gene flow, IM, SC, AM, “hm’” models). Here we considered two categories of loci: barrier loci without migration (*m* = 0) and loci for which *m* is freely estimated (m > 0). We modeled heterogeneity in effective population size in all four models (“hn” models) allowing for evolution of two sets of loci with different effective population sizes (*N_e_*_1_ and *N_e_*_2_). We further simulated scenarios including heterogeneity in both migration rate and population size (“hmhn” models) with four sets of loci: m = 0 and *N_e_*_1_, m = 0 and *N_e_*_2_, m > 0 and *N_e_*_1_, and m > 0 and *N_e_*_2_. In total, we assessed 18 demographic models and optimized each with the observed joint SFS using 20 replicate runs with perturbed parameters as starting points. We excluded SC models that did not converge and had very short periods of unrealistically high gene flow (2*N_e_m* > 30) (personal communication, Ryan Gutenkunst). Nested model comparisons were performed using likelihood ratio tests; comparisons of non-nested models were based on the Akaike Information Criterion (AIC).

We estimated the demographic parameters from the best-support model, including divergence time, population sizes, migration rates, and proportions of the different types of loci (where included), with 95% confidence intervals constructed using 500 rounds of re-sampling SNPs from contigs. In ∂a∂i, divergence time is inferred in units of 2*N_ref_* generations. *N_ref_* constitutes the ancestral effective population size and can be estimated based on *θ = 4 N_ref_ µ L*, where *θ* denotes the population mutation rate, *µ* the mutation rate per site per generation, and *L* the effective sequence length involved. We estimated *L* as the total base number of the reference contigs meeting filtering criteria. We used a generation time of one year and a mutation rate *µ* of 7.92 × 10^-9^ (Krasovec et al., 2018).

### Calculation of differentiation, sequence divergence and diversity statistics

We calculated differentiation between the two species as hierarchical *F*-statistics (“hierarchical *F*_ST_”) to account for population structure within species using AMOVA models in the R package *hierfstat* (Goudet, 2004) for both SNPs and contigs. We also calculated Neís estimate of *F*_ST_ between all population pairs (between species and within species). We calculated sequence divergence, *d*_XY,_ between the two species using a custom Perl script. Within each species, we computed genetic diversity (π) using VCFtools (Danecek et al., 2011) and estimated Tajima’s D using a custom Perl script. For contig-based calculations dependent of sequence length (*d*_XY_, π and Tajima’s D), we determined contig length as number of bases meeting SNP filtering criteria (read depth > 5 per individual; coverage: >70% of the individuals).

### Genome scans using linkage maps

We examined genome-wide patterns in differentiation, sequence divergence and diversity statistics by mapping the contigs onto the two linkage maps constructed in a previous study from two F_2_ crosses between *S. dioica* and *S. latifolia*: F_2DL_ (1470 markers) and F_2LD_ (1265 markers) (Liu & Karrenberg, 2018). The two linkage maps are treated separately here because only 20% of the contigs with mapped markers are shared between the two maps (Liu & Karrenberg, 2018). Uncertainty in marker order on a consensus map, particularly in marker-dense regions, may lead to spurious patterns. We used the local polynomial regression (LOESS) curve along each linkage group to represent patterns for each statistic (Cleveland, 1979).

### Genomic landscapes and signatures of selection

As a first step, we analyzed correlations between contig-based *F*_ST_ and *d*_XY_ using a permutation test in the R package *coin* (Hothorn, Hornik, Wiel, & Zeileis, 2006). We further identified genomic islands of elevated differentiation as contig sequences with hierarchical *F*_ST_ exceeding the 95% quantile of the overall *F*_ST_ value distribution. To assess signatures of selection in these highly differentiated regions, we compared *d*_XY_, π and Tajima’s D between the differentiation islands and background regions using Mood’s median test via the R package *coin* (Hothorn et al., 2006). The above analyses were performed both for mapped contigs (data from both linkage maps combined) and for all contigs in our dataset.

### Within vs. between species differentiation

Genetic differentiation landscapes within and between species are expected to correlate if they are mainly governed by background selection and conserved genomic features (Burri, 2017; Vijay et al., 2017). We assessed the role of background selection for the generation of the differentiation landscape by comparing differentiation landscape between species (hierarchical *F*_ST_) to differentiation landscapes within species using the population pairs with the highest *F*_ST_ within each species. We further evaluated correlations (across contigs) of hierarchical *F*_ST_ and within-species *F*_ST_ for both mapped and all contigs using a permutation test the R package *coin* (Hothorn et al., 2006).

### Differentiation near QTLs for reproductive barrier traits

We first compared the location of highly differentiated regions to the previously identified QTLs for traits associated with reproductive barriers between *Silene dioica* and *S. latifolia* (Liu & Karrenberg, 2018): assortative mating (flower color, flower size, flower number, stem height), habitat adaptation (first-year flowering, specific leaf area, leaf succulence, survival), phenology (flowering time) and hybrid male fertility reduction (pollen number). As a second step, we assessed whether genetic differentiation in regions associated with different reproductive barriers (contigs within 1 cM of QTLs) had significantly higher medians of hierarchical *F*_ST_ between species than or all mapped markers using a randomization test. We obtained the distributions of medians under the null hypothesis (no difference) using of 10, 000 sets of randomly selected contigs of the same size as the number of contigs associated each barrier type. We determined the percentile of the observed medians in the respective distributions of medians from randomly selected datasets and expressed P-values as 1 minus this percentile.

### Patterns of isolation by distance within and between species

We examined the relationships between pair-wise *F*_ST_ between populations (within *Silene dioica*, within *S. latifolia* and between species) and geographical distance for differentiation islands and for regions of low differentiation (hierarchical *F*_ST_ below the 25% quantile of the overall distribution). Correlation tests between *F*_ST_ and geographic distance were performed using Mantel permutation tests implemented in the R package *vegan* (Oksanen et al., 2019).

## Results

### Sequencing output

We obtained on average 4,303,869 reads per individuals from ddRAD sequencing, which in total yielded 1,799,882,123 reads. After filtering, we retained 87,006 SNPs for the downstream analysis with an average read coverage of 21.2. These SNPs were located on 10,415 contigs with an average length of 251 bp. 1327 contigs could were present on linkage maps (817 contigs on the F_2DL_ map, 632 contigs on the F_2LD_ map, 122 contigs on both maps).

### Population structure and overall hierarchical F_ST_

We used a set of 36,413 SNPs after pruning for linkage disequilibrium to analyze population structure. With *K* = 2, two groups are clearly separated as well, corresponding to *S. dioica* and *S. latifolia*, with limited admixture in a few populations (Figure 1b), particularly especially, the Swiss populations of both species (CH1 and CH3) and the Belgian population of *S. latifolia* (BEL1). With *K* = 20, the number of natural populations included in the study, all populations could be recognized, although there was evidence for considerable introgression within species (Figure S3, supporting Information). A phylogenetic tree based on genetic distance also showed two clusters, consisting of populations from each species (Figure 1c).

The overall hierarchical *F*_ST_ between *S. dioica* and *S. latifolia* was 0.28, as estimated from AMOVA analysis (Table 1). Average pair-wise weighted *F*_ST_ was 0.357 (range: 0.261 – 0.424) for between species population pairs, 0.099 (range: 0.019 – 0.165) for population pairs within *S. dioica* and 0.136 (range: 0.027 - 0.267) for population pairs within *S. latifolia* (Fig. 1d). This is in agreement with the phylogentic tree, where branch lengths of *S. latifolia* populations were longer than those of *S. dioica* populations (Figure 1c).

**Table 1.** Summary of population genetic statistics for the campions *Silene dioica* (11 populations) and *S. latifolia* (9 populations) based on reduced-representation sequencing (ddRAD-seq).

| Statistic | Value |  |
| --- | --- | --- |
|  | Mean | Median |
| Number of SNPs | 87006 |  |
| Number of contigs | 10415 |  |
| hierarchical $F_{ST}$ between species | 0.278 | |
| $d_{XY}$ between species | $5.28 \times 10^{-3}$ | $3.36 \times 10^{-3}$ |
| pair-wise $F_{ST}$ within *S. dioica* | 0.098 | 0.096 |
| pair-wise $F_{ST}$ within *S. latifolia* | 0.136 | 0.128 |
| $\pi$ within *S. dioica* | $2.18 \times 10^{-3}$ | $1.40 \times 10^{-3}$ |
| $\pi$ within *S. latifolia* | $2.48 \times 10^{-3}$ | $1.70 \times 10^{-3}$ |
| Tajima's D within *S. dioica* | -0.156 | -0.475 |
| Tajima's D within *S. latifolia* | -0.156 | -0.435 |

### Demographic history of lineage divergence

Models involving migration generally outperformed the strict isolation (SI) models significantly, based on AIC (Figure 2, Figure S2, Table S2, Supporting Information). These models, isolation with migration (IM), secondary contact (SC) and ancient migration (AM), were slightly improved by adding population expansion (“exp” models) but substantially improved by adding heterogeneity in migration rate (“hm” models), effective population size (“hn” models) or both migration rate and population size (“hmhn” models, Figure 2, Table S2, Supporting Information). Under each scenario (IM, SC and AM), the model including heterogeneity in both population size and migration rate (“hmhn” models) had the best fit, as indicated by likelihood ratio tests (Table S2, Supporting Information). The overall best-supported model in terms of AIC was the isolation with migration model with heterogeneous population size and migration rate (“IMhmhn”, Figure 2, Table S2, Supporting Information). SC and AM models with heterogeneous population sizes and migration rates (“SChmhn”, “AMhmhn”) fit the data nearly as well; however, in both scenarios, the estimated duration of the phase without gene flow was close to zero, such that they converge to the IMhmhn model (Tables S2 and S3, Supporting Information).

**Figure 2.**
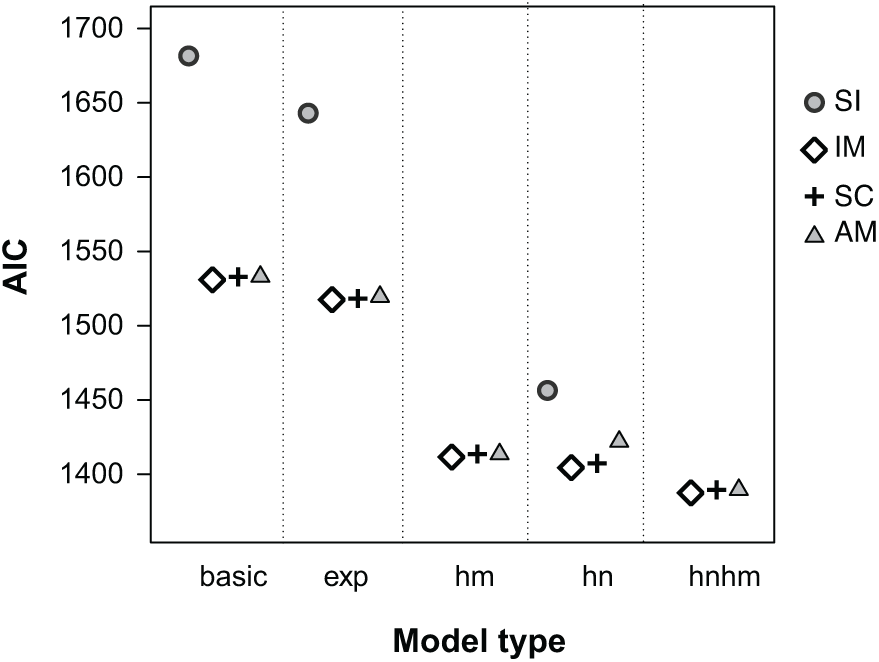
Akaike’s information criterion (AIC) for demographic models of the lineage split between the campions *Silene dioica* and *S. latifolia* in four scenarios: strict isolation (SI), isolation with migration (IM), secondary contact (SC) and ancient migration (AM). Models contained no further parameters (“basic”), allowed for populations expansion (“exp”), or included heterogeneity in migration rate (“hm”), effective population size (“hn”) or both parameters (“hmhn”). “hm” and “hmhn” models were not calculated for the SI scenario which does not include any migration.

The best-supported IMhmhn model reflected the joint site frequency spectrum fairly well and residuals were mainly in the range of −2 to 2 (Figure 3, Figure S3, Supporting Information). Under this model, 39% of the loci had reduced effective population size and 5% of the loci conformed to a scenario of divergence without gene flow (Table 2). Most of these potential barrier loci had reduced *N_e_* (4.4% of all loci with *m* = 0 and low *N_e_* and *m* = 0; 0.4% of all loci with *m* = 0 and high *N_e_*). It is worth noting that adding heterogeneity in both *m* and *N_e_* strongly affected estimates of the proportion of loci without gene flow.

**Figure 3.**
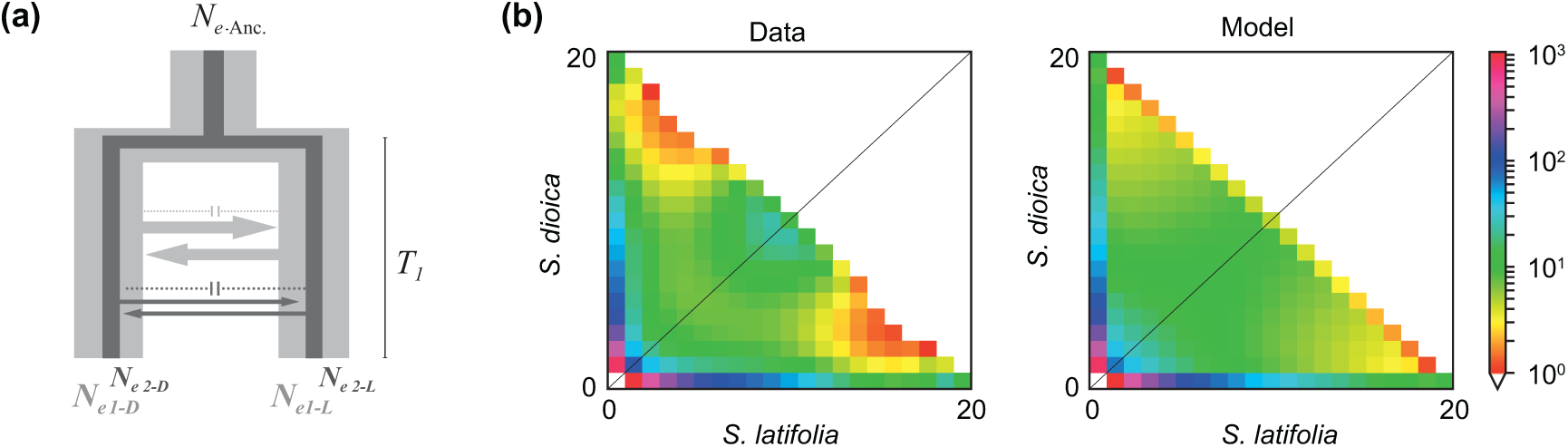
**(a)** Illustration of the best-supported demographic model, isolation with migration with heterogeneous migration rate and population size (“IMhmhn”). Light grey represents loci evolving with large population size (*N_eD1,_ N_eL1_*) and dark grey represents the loci with reduced population size (*N_eD2,_ N_eL2_*); arrows indicate migration and barrier loci (dotted lines, not drawn to scale). **(b)** Joint site frequency spectra of the data and under the IMhmhn model.

**Table 2.**
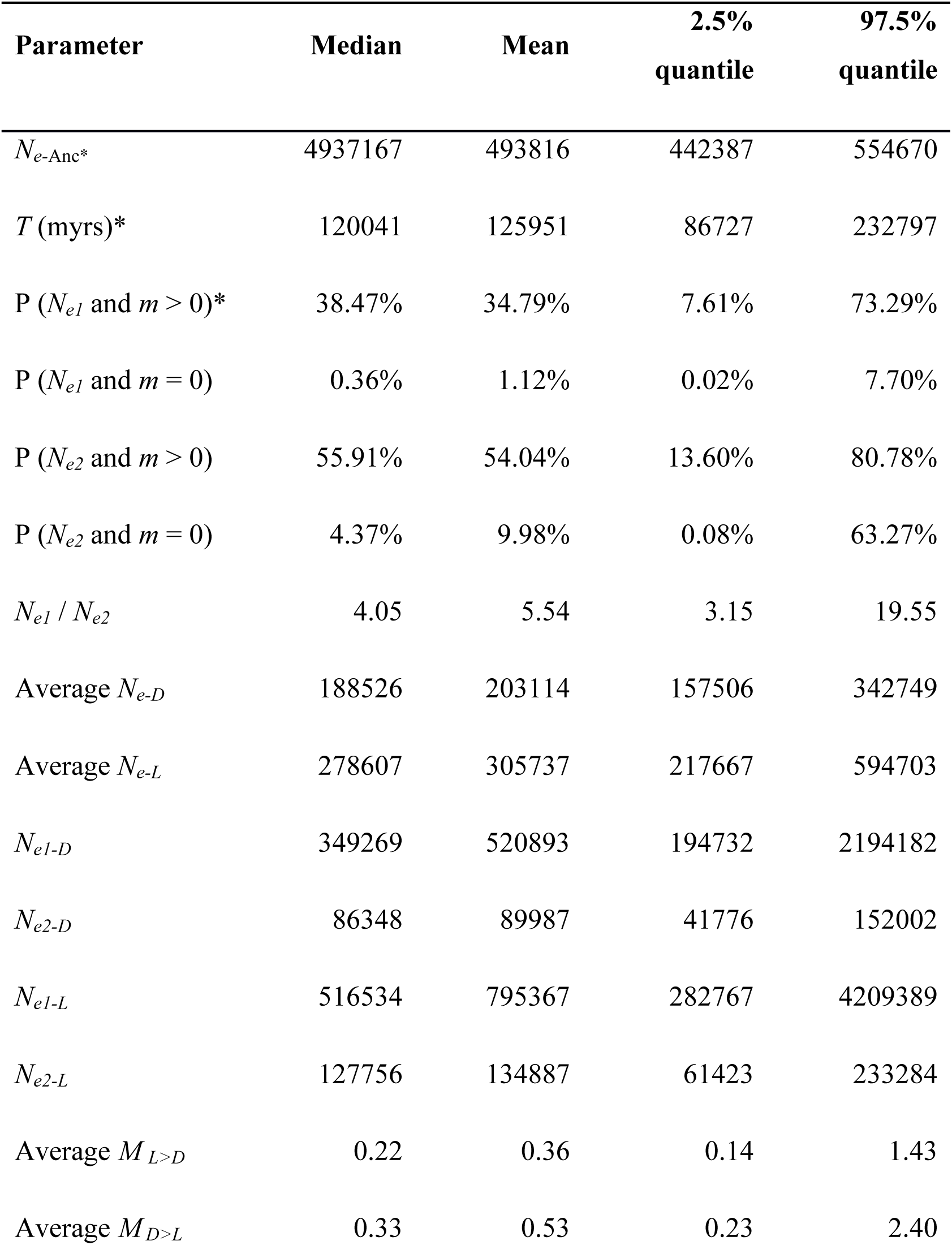
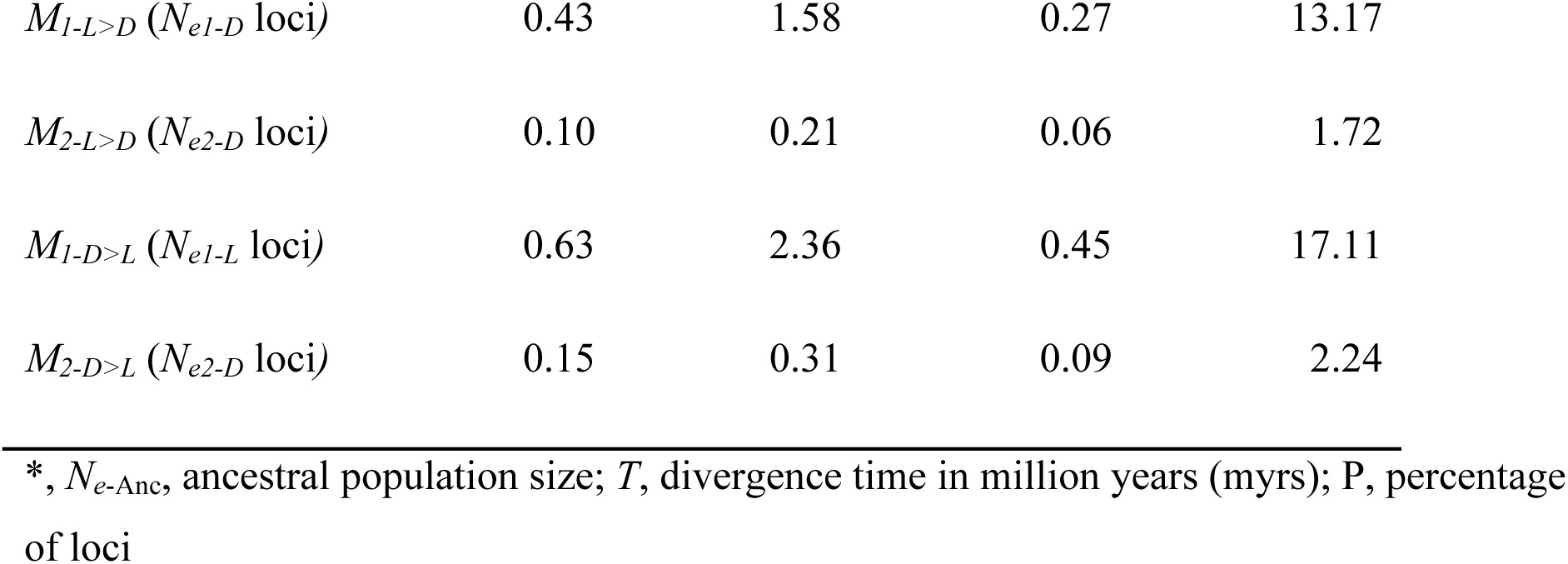
Demographic parameters in the best-supported demographic model (IMhmhn) of the lineage divergence between the campions *Silene dioica* and *S. latifolia*: isolation with migration with heterogeneous migration rate (*m* = 0 and *m* > 0) and heterogeneous effective population size (*N_e1_* and *N_e2_*); subscripts stand for *S. dioica* (D) and *S. latifolia* (L) and for the direction of gene flow, from *S. dioica* into *S. latifolia* (D> L) and vice versa (L>D).

The estimated median population migration rate across the genome from the IMhmhn model was 0.22 (95% CI: [0.14-1.43]) migrants per generation from *S. latifolia* to *S. dioica* and 0.33 (95% CI: [0.23-2.4]) from *S. dioica* to *S. latifolia* (Table 2). We obtained an effective population size of 188,526 (95% CI: [157,506-342278]) for *S. dioica* and 278,607 (95% CI: [217,667-594,702]) for *S. latifolia*. The divergence time between *S. dioica* and *S. latifolia* was estimated to be 0.12 million years (95% CI: [0.087-0.23] myrs) assuming a generation time of one year (Table 2).

### Genomic landscape of differentiation, sequence divergence and diversity

Hierarchical *F*_ST_ on both SNP- and contig-based levels fluctuated widely across the genome (Figure 4, Figure S4, Supporting Information). Both hierarchical *F*_ST_ and sequence divergence, *d*_XY_, reached peaks around the middle of most linkage groups on both linkage maps (Figure 4, Figure S4, Supporting Information), in spite of largely different SNP markers (Liu & Karrenberg, 2018). *F*_ST_ and *d*_XY_ were significantly correlated (mapped contigs: r = 0.347, *P* < 0.001, all contigs: r = 0.464, *P* < 0.001, based on 9999 permutations). Highly divergent regions in the middle of linkage groups often contained loci with reduced nucleotide diversity (π) and more negative Tajimás D in one or both species (Figure 4, Figures S4, S5, S6, Supporting Information).

**Figure 4.**
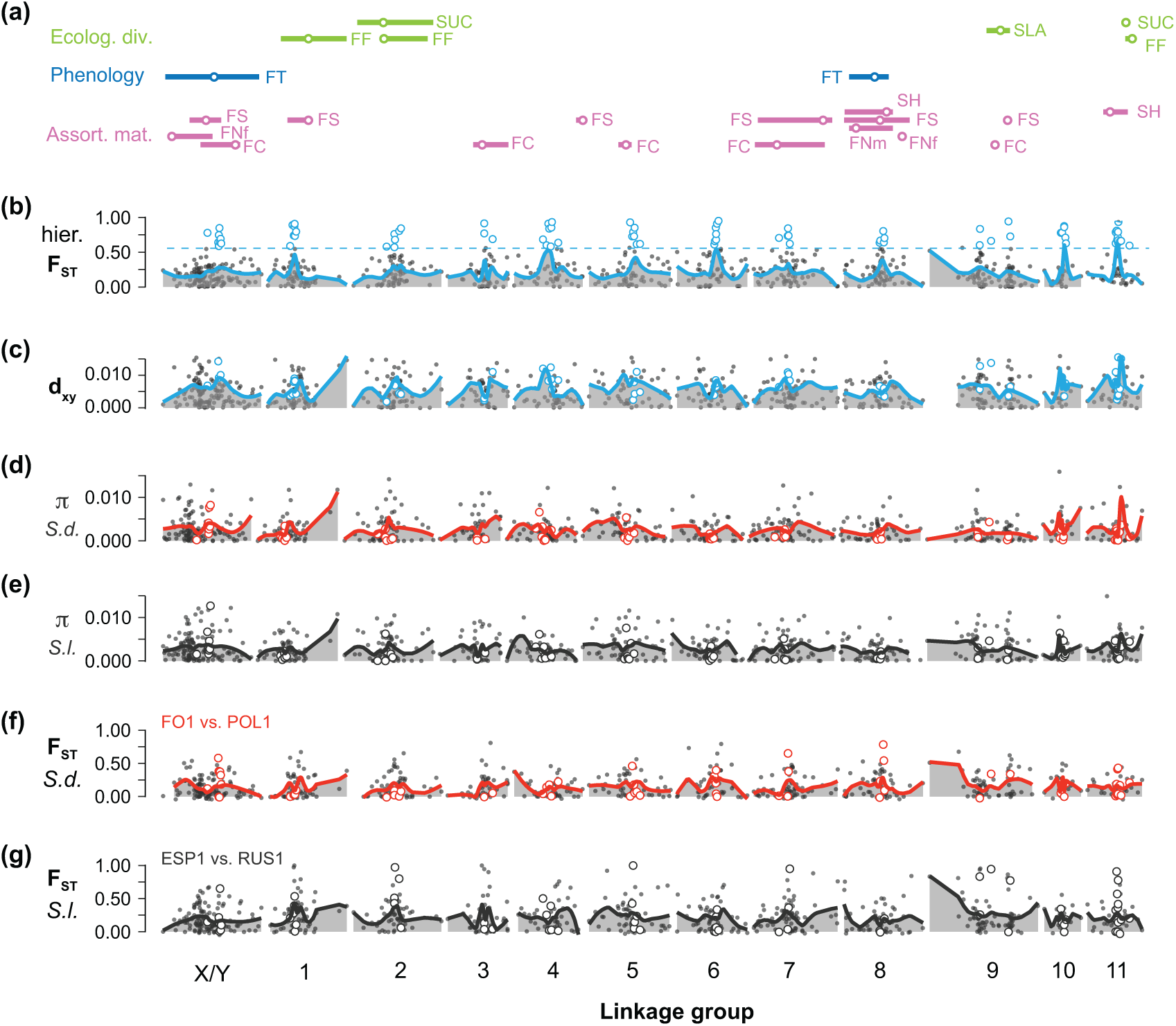
Genomic landscapes and QTLs for traits associated with reproductive barriers between the campions *Silene dioica* and *S. latifolia* based on a linkage map of the F_2DL_ cross (for the F_2LD_ cross see Figure S3, Supporting Information); **(a)**, locations (circles) and 1.5 LOD drop intervals (lines) of QTLs related ecological divergence (FF: first-year flowering, SUC: leaf succulence, SLA: specific leaf area), phenology (FT: flowering time) and assortative mating (FC: flower color, FS: flower size, FNf and FNm: flower number for females and for males, SH: stem height); **(b)**, hierarchical *F*_ST_ between the two species with the 95% quantile of overall *F*_ST_ distribution (dashed line); **(c)**, sequence divergence (*d*_XY_); **(d)** and **(e)**, nucleotide diversity (π) within *S. dioica* (*S.d.*) and within *S. latifolia* (*S.l.*); **(f)** and **(g)**, *F*_ST_ of the most differentiated population pair within *S. dioica* and within *S. latifolia.* Lines in panels **(b)** - **(g)** are drawn using LOESS functions; empty circles represent contigs with hierarchical *F*_ST_ exceeding the 95% quantile (genomic islands of differentiation).

For mapped contigs, *d*_XY_ was significantly elevated in genomic islands of differentiation as compared to the genomic background, and both π and Tajima’s D were significantly reduced in both species (Figure 5, Table S4, Supporting Information). Results from all contigs were very similar, except that π was not significantly reduced in differentiation islands within *S. latifolia* (Figure S7, Supporting Information, Table S4, Supporting Information).

**Figure 5.**
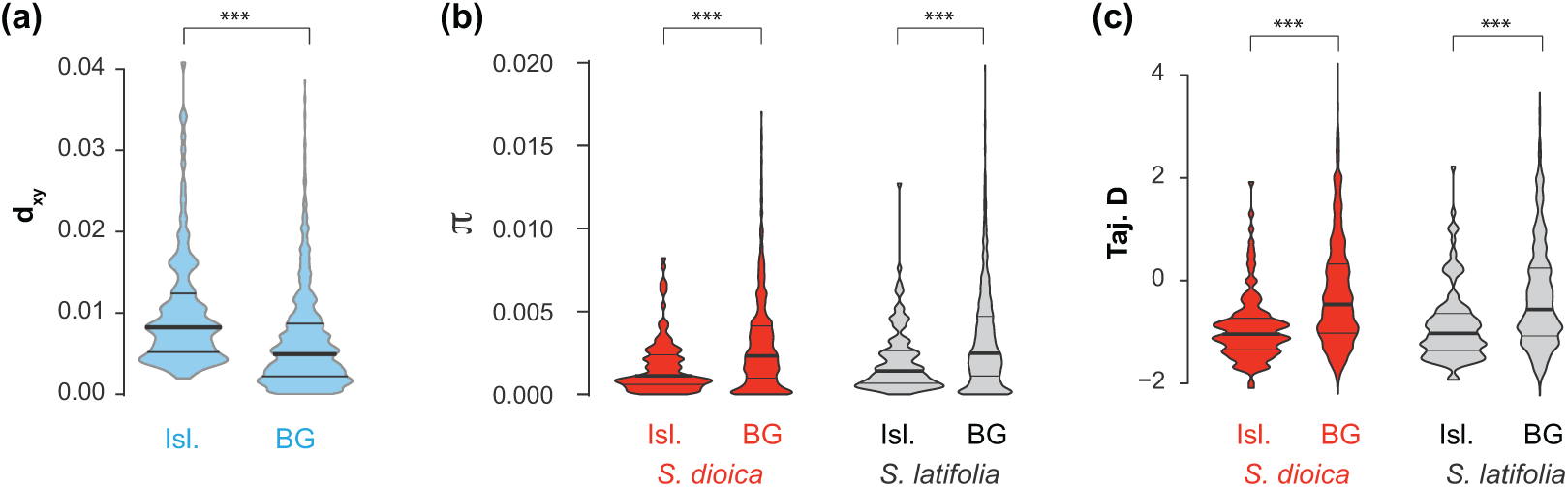
Violin plots for population genetic statistics in the campions *Silene dioica* and *S. latifolia* in genomic islands of differentiation (Isl.) and the genomic background (BG) for contigs mapped onto linkage maps: **(a)** sequence divergence (*d*_XY_); **(b)** genetic diversity (π), and **(c)** Tajima’s D. Within each violin, the horizontal lines represent the 25%, 50% (median, thick lines) and 75% quantiles. Results from Mood’s median tests are given on the top (***, P-Value < 0.001).

### Within- vs. between-species differentiation

The genomic landscapes of differentiation between species generally did not coincide with that of highly differentiated population pairs within *S. latifolia* (ESP1 vs. RUS1, overall *F*_ST_ = 0.27) or within *S. dioica* (FO1 vs. POL1, overall *F*_ST_ = 0.16, Figure 4). Within species comparisons had less pronounced *F*_ST_ peaks in the middle of several linkage groups as compared to between species hierarchical *F*_ST_, whereas the remaining linkage groups showed no obvious differentiation peaks in within species comparisons (Figure 5, Figure S4, Supporting Information). Correlations of within species *F*_ST_ and between species hierarchical *F*_ST_ were weak (mapped contigs: r = 0.028, P = 0.319 for *S. dioica* and r = 0.11, P < 0.001 for *S. latifolia*; all contigs: r = 0.071, P < 0.001 for *S. dioica* and r = 0.147, P < 0.001 for *S. latifolia*, Figure S8, Supporting Information).

### Differentiation near QTLs for reproductive barrier traits

QTLs for traits associated with reproductive barriers often resided in genomic regions with high *F*_ST_ and *d*_XY_ were concentrated (Figure 4, Figure S4, Supporting Information). Contigs mapping within 1 cM of QTLs associated with assortative mating had significantly elevated median *F*_ST_ (0.249) as compared to the median *F*_ST_ of all mapped contigs (0.192, P = 0.039, randomization test); but this was not true for contigs near QTLs associated with habitat adaptation (median *F*_ST_: 0.098, P = 0.992), phenology (median *F*_ST_: 0.184, P = 0.564) or hybrid male fertility reduction median *F*_ST_: 0.275, P = 0.097, Figure 6).

**Figure 6.**
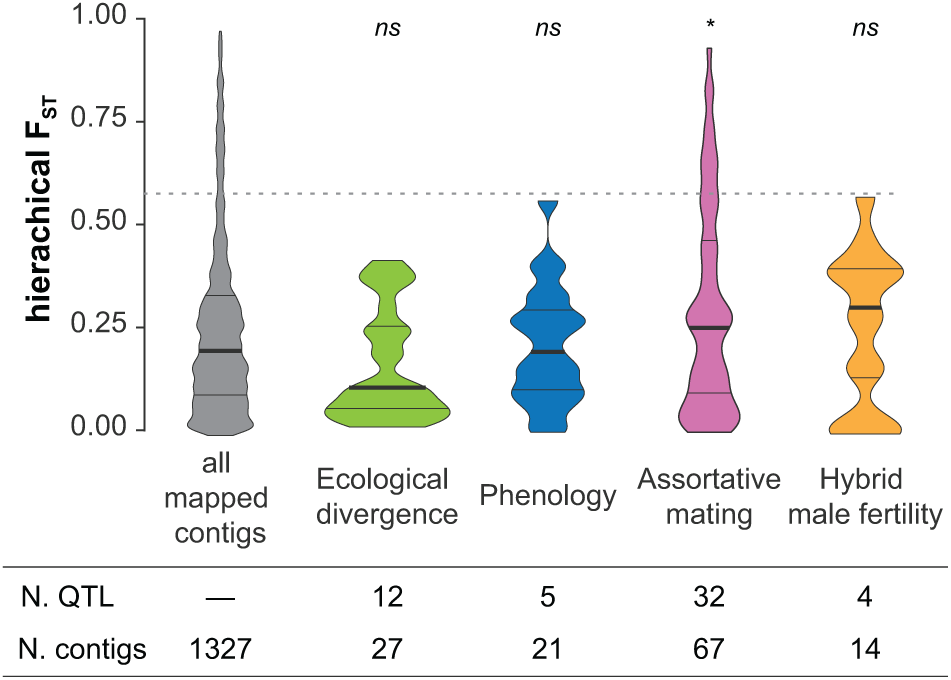
Violin plot of *F*_ST_ values for all contigs mapped onto linkage maps and for contigs within 1 cM of QTLs related to ecological divergence, phenology, assortative mating and male fertility (only detected in cross F_2LD_ cross see Figure S3, Supporting Information). Numbers of QTLs and contigs in each category are given below each violin. Results of randomization tests for elevations of median *F*_ST_ in QTL-associated contigs as compared to all contigs are given at the top (*ns*, P > 0.05; *, P < 0.05).

### Patterns of isolation by distance

Within *S. dioica*, pair-wise *F*_ST_ was only weakly associated with geographical distances (Figure 7a, Figure S9a, Table S5, Supporting Information). Within *S. latifolia*, in contrast, we generally detected significant associations of geographic distance with pairwise *F*_ST_ isolation by distance patterns (Figure 7b, Figure S9b, Table S5, Supporting Information). In both species, differentiation islands exhibited slightly elevated pair-wise *F*_ST_ and a tendency for a weaker correlation between pair-wise *F*_ST_ and geographic distance as compared to loci with low between-species differentiation (Figures 7, Figure S9, Table S5, Supporting Information). In between-species population pairs, pair-wise *F*_ST_ correlated strongly and significantly with geographical distance at loci with low between-species differentiation. At differentiation islands, in contrast, no significant increase of pair-wise *F*_ST_ with geographical distance was detected, suggesting that many of these loci constitute barrier loci (Figure 7c, Figure S9c, Table S5, Supporting Information). For these latter analyses we excluded pair-wise comparisons to one *S. latifolia* population (CH2) that showed strongly reduced pair-wise *F*_ST_ for highly differentiated regions, and moderately reduced pair-wise *F*_ST_ for regions with low differentiation (Figure 7, Figure S9). This population also exhibited signs of admixture (Figure 1b).

**Figure 7.**
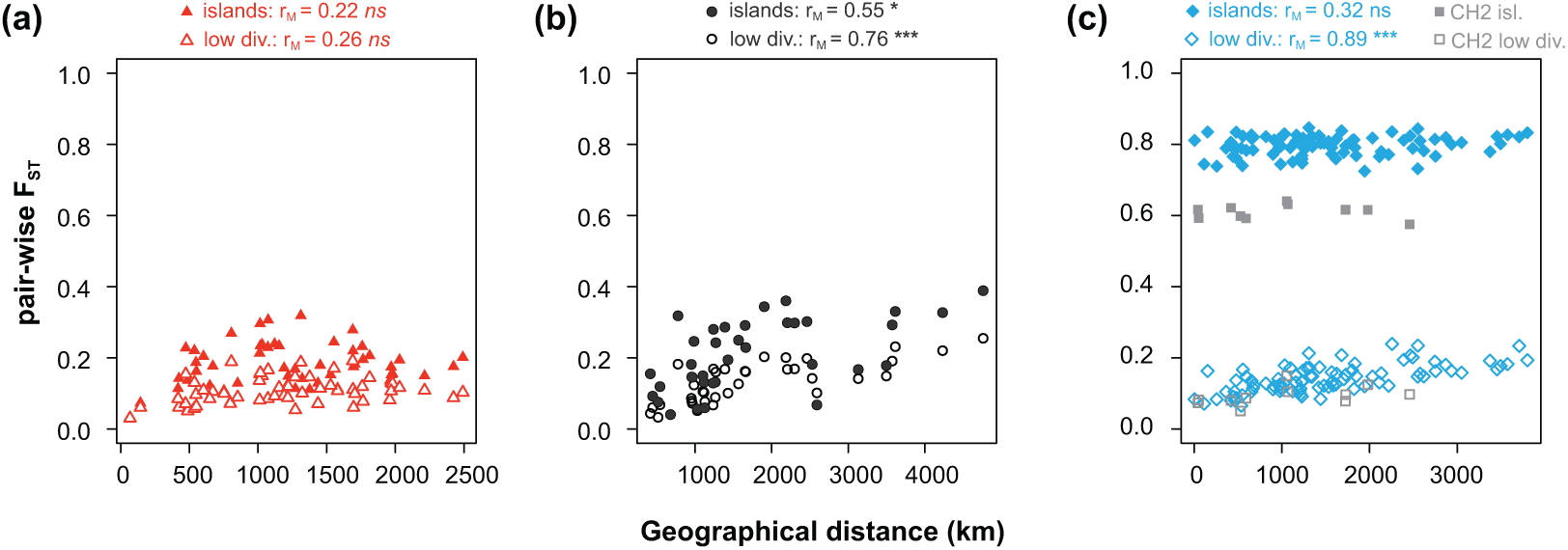
Scatterplots of pair-wise differentiation (*F*_ST_) against geographic distance (km) for population pairs within the campion *Silene dioica* **(a)**, within *S. latifolia* **(b)**, and between the two species **(c)** based on contigs mapped to linkage maps. Pair-wise *F*_ST_ is given for between-species differentiation islands (filled symbols(> 95% quantile) and for regions of low between-species divergence (empty symbols, < 25% quantile) based on mapped contigs. Results of Mantel tests are indicates at the top; *ns*, P > 0.05; *, P < 0.05; ***, P < 0.001.

## Discussion

In this study, we investigated the evolutionary processes leading to divergence between two closely related and hybridizing campion species, *Silene dioica* and *S. latifolia*. We sampled populations throughout the distribution ranges of each species and generated reduced representation sequencing data. We used an integrated approach and performed demographic modeling combined with population genomic analyses. Together, our results support a model where the two species diverged with continuous gene flow but have evolved barrier loci at which gene flow is reduced. Our analyses further indicate that divergent selection contributed to the evolution of putative barrier loci while background selection played a limited role.

### Demographic history supports divergence with gene flow and barrier loci

Demographic analyses suggest that *S. dioica* and *S. latifolia* diverged with continuous gene flow, in line with evidence for ongoing hybridization in natural populations of the two species (Karrenberg & Favre, 2008; Minder et al., 2007). Similar scenarios of comparatively advanced divergence (hierarchical *F*_ST_ = 0.28 in our study) combined with persistent gene flow are only rarely reported, for example in Japanese sticklebacks (Ravinet et al., 2018) and *Heliconius* butterflies (Martin et al., 2013). Demographic analyses in many other systems, in contrast, point to divergence in para/allopatry with gene flow only during secondary or intermittent contact (Christe et al., 2017; Duranton et al., 2018; Tine et al., 2014). In our analysis, secondary contact (SC) and ancient migration (AM) models had only slightly less support than the best-supported isolation with migration (IM) models (but with an AIC difference exceeding 2). However, periods without gene flow in AM and SC models were extremely short, such that they effectively converged to the best-supported IM model. Nonetheless, we cannot exclude more complex demographic scenarios such as multiple short periods of allopatry combined with population expansion (Christe et al., 2017). Large effects of population expansion are unlikely though, based on simple IM models with population expansion in this study, as well as in (Muir et al., 2012) and in Guirao-Rico et al. (2017). We are therefore confident that our isolation with migration model captures the main aspects of the two species’ demographic history.

Our demographic models included and supported heterogeneity in both migration rate and effective population size. This is in line with results from various animal systems where heterogeneous migration rates were most common at intermediate stages of speciation, whereas support for heterogeneity in effective population size varied across systems irrespective of the speciation stage (Roux et al., 2016). We estimated that 5% of the loci evolved without gene flow and thus potentially barrier loci. Our analyses further indicate that these putative barrier loci were concentrated in regions with reduced effective population sizes as predicted by models of speciation with gene flow (Ortiz-Barrientos & James, 2017; Ravinet et al., 2017; Seehausen et al., 2014; Yeaman et al., 2016). However, effects of heterogeneity in population size and migration on the joint site frequency spectrum can be difficult to distinguish (Christe et al., 2017; Rougemont & Bernatchez, 2018; Roux et al., 2016).

Our results generally agree with previous demographic analyses in the two species that used more limited datasets and simpler models but also rejected strict isolation scenarios in favor of divergence with gene flow (Guirao-Rico et al., 2017; Hu & Filatov, 2015; Muir et al., 2012). We estimated divergence time to 0.12 million years before present, considerably earlier than estimates of Muir et al. (2012), Hu and Filatov (2015) and Guirao-Rico et al. (2017). This is likely a consequence of including heterogeneity in population size and migration rate in demographic models in this study, as indicated by comparisons of estimates under the best-supported model and under a model similar to those employed in previous studies (Guirao-Rico et al., 2017; Hu & Filatov, 2015; Muir et al., 2012). Our divergence time estimate places the lineage split close to the onset of the most recent glaciation in Europe with rapid environmental changes (Hewitt, 2000). However, this estimate is contingent on a generation time of one year observed for these species (Favre et al., 2017; Hu & Filatov, 2015; Muir et al., 2012) but generation time can also be two or three years in colder environments (Favre et al., 2017). Average migration rate estimates (*M =* 2*N_e_m*) from our best-supported model (0.22 [CI: 0.14 - 1.43] and 0.33 [0.23 - 2.4] from *S. latifolia* into *S. dioica* and vice versa) were similar to those reported by Muir et al. (2012) but lower than those reported by Hu and Filatov (2015). Estimates of migration rate were further consistent with near-complete reproductive isolation, mostly through adaptation to the habitat and to pollinators (Karrenberg et al., 2019). The proportion of the gene pool replaced by the other species per generation *m*, was estimated to ca. 6 x 10^-7^ here, and can be compared for the probability of F_1_ hybrid production, P_hyb_ (Sambatti, Strasburg, Ortiz-Barrientos, Baack, & Rieseberg, 2012). *m* was several orders of magnitude smaller than P_hyb_ estimated from the strength of reproductive isolation (P_hyb_ = 1-RI_total_: 0.010-0.036, depending on the direction), suggesting that additional ecological or genetic reproductive barriers may exist (Karrenberg et al., 2019). Overall, we thus establish realistic demographic estimates for a likely case of speciation in the face of gene flow in plants.

### Accentuated differentiation (F_ST_) and sequence divergence (d_XY_) in linkage group centres suggest the evolution of barrier loci

Genetic differentiation, in terms of hierarchical *F*_ST_ between species, and sequence divergence (*d*_XY_) were both accentuated in the middle of most linkage groups. Elevated differentiation in chromosome centers has been described for diverse taxa and attributed to reduced recombination rates in the middle of metacentric chromosomes (Berner & Roesti, 2017; Nachman & Payseur, 2012). Chromosomes of *S. latifolia* are indeed metacentric or sub-metacentric (Lengerova et al., 2004); however, direct recombination rate estimates are not available for *Silene* to date. In our study, *F*_ST_ and *d*_XY_ were positively correlated and highly differentiated regions (genomic islands of differentiation) had an approximately two-fold higher median *d*_XY_ as compared to remaining genomic regions. Moreover, linkage group centers often harbored previously identified QTLs for reproductive barriers traits (see also below) with evidence of clustering (Liu & Karrenberg, 2018). In a divergence-with-gene flow scenario, as in the species studied here, these results are suggestive of reductions in gene flow and thus barrier loci co-localizing in low-recombination regions (Cruickshank & Hahn, 2014; Nachman & Payseur, 2012; Rafajlović, Emanuelsson, Johannesson, Butlin, & Mehlig, 2016; Ravinet et al., 2017; Seehausen et al., 2014; Yeaman et al., 2016); however, we cannot exclude that lineage sorting in the ancestor also contributed to the pattern (Pease, Haak, Hahn, & Moyle, 2016; Richards et al., 2019). Evidence for genomic islands with elevated *F*_ST_ and *d*_XY_ are comparatively rare and have been reported for Darwińs finches (Han et al., 2017) European sea bass (Duranton et al., 2018) and poplars (Wang et al., 2016). Our findings are in stark contrast to many other systems where differentiation landscapes are mainly shaped by background selection acting on a heterogeneous recombination landscapes: under this scenario, *d*_XY_ is not elevated or even reduced in regions of high differentiation and differentiation landscapes across replicate species pairs are correlated (Burri, 2017; Nachman & Payseur, 2012), as reported in sunflowers (Renaut et al., 2013) and different bird lineages (Burri et al., 2015; Delmore et al., 2018; Vijay et al., 2017). In our study, we found only weak correlations of *F*_ST_ between species with *F*_ST_ between highly divergent populations within species, and *F*_ST_ between within-species population pairs was only slightly elevated at between-species islands of differentiation. This may indicate limited effects of background selection; however, loci diverging with gene flow may also repeatedly arise in the same regions, for example, low-recombination regions (Berner & Roesti, 2017; Samuk et al., 2017; Yeaman et al., 2016). Our results are thus consistent with and important role of barrier loci in generating the differentiation landscape.

### Signatures of selection at highly differentiated regions

Genomic islands of high differentiation show significant signatures of selection (Nielsen, 2005) within both species: reduced sequence diversity and an overrepresentation of rare variants (reduced and negative Tajima’s D). Together with our results on sequence divergence, *d*_XY_, these findings suggest that highly differentiated regions, containing potential barrier loci, arose through divergent selection. Reproductive isolation between the two species is mainly associated with adaptation to the habitat and, presumably, to pollinators (Favre et al., 2017; Goulson & Jerrim, 1997; Karrenberg et al., 2019). Signatures of selection in highly divergent regions in other systems have been detected in ecotypes or early-stage ecological speciation in three-spined sticklebacks (Feulner et al., 2015; Marques et al., 2017; Samuk et al., 2017) and in cichlids (Malinsky et al., 2015; Meier et al., 2018), as well as in more advanced stages of speciation in poplars (Wang et al., 2016). Our results thus provide additional evidence for an important role of divergent selection in generating the genomic landscape of differentiation.

### Elevated differentiation near QTLs for reproductive isolation traits

We detected elevated differentiations in 1 cM regions surrounding previously identified QTLs (Liu & Karrenberg, 2018) associated with assortative mating but not for QTLs associated with other reproductive barriers (Karrenberg et al., 2019) such as habitat adaptation, phenology or hybrid male fertility reduction. This could be due to a particularly important role of assortative mating as a reproductive barrier (Coyne & Orr, 2004; Marques et al., 2017; Martin et al., 2013; Riesch et al., 2017). Alternatively, this finding may result from a simpler genetic architecture of floral traits as compared ecological divergence - adaptive divergence in traits with complex genetic architectures may not lead to high differentiation at any of the involved loci (Liu & Karrenberg, 2018; Rockman, 2012; Yeaman, 2015). However, these patterns are very difficult to interpret, given the scarce genome coverage of the reduced representation sequencing data used here and the large confidence intervals of some the QTL location. Nonetheless, our results are consistent with our interpretation that adaptive divergence and the ensuing reproductive barriers shaped the genomic landscape of differentiation in this system.

### Highly differentiated regions are barriers to gene flow between species

Within *S. latifolia*, we detected significant isolation by distance patterns consistent with post-glacial range expansion, whereas an isolation by distance pattern was very weak *S. dioica*, possibly because geographical distance does not reflect dispersal routes very well in mountainous regions (Hathaway, Malm, & Prentice, 2009; Prentice, Malm, & Hathaway, 2008; Rautenberg, Hathaway, Oxelman, & Prentice, 2010). In between-species population pairs, *F*_ST_ significantly increased with geographical distance at loci with low differentiation between species, indicating that the two species are connected by gene flow. At highly differentiated loci, in contrast, no significant *F*_ST_ increase with geographical distance was detected, suggesting that many of these loci constitute barrier loci. This is in line with results of heterogeneous gene flow from our demographic analysis. Between-species comparisons involving one *S. latifolia* population from Switzerland, with comparatively high admixture, had generally reduced between-species *F*_ST,_ particularly for highly differentiated loci. This is likely due to recent hybridization and introgression in this population. Ongoing hybridization has previously been documented for this area (Karrenberg & Favre, 2008; Minder et al., 2007). Overall, these results suggest that highly differentiated regions correspond to between-species barriers to gene flow.

### Conclusion

Multiple lines of evidence support the evolution of barrier loci during speciation with gene flow in the campions *Silene dioica* and *S. latifolia*: (1) demographic analyses indicate divergence with continuous and heterogeneous gene flow, (2) differentiation (*F*_ST_) and sequence divergence (*d*_XY_) were positively correlated and elevated in the middle of most linkage groups (3) highly differentiated regions exhibited signatures of selection, and differentiation was elevated near QTLs for reproductive barrier traits, and (4) isolation by distance patterns suggest that *S. dioica* and *S. latifolia* are connected by gene flow at regions with low differentiation but not at islands of high differentiation. Strong cumulative reproductive isolation in this system results mainly from adaptation to the habitat and to pollinators (Karrenberg et al., 2019) with polygenic and clustered genetic architectures (Liu & Karrenberg, 2018). The results of this study thus align well with theoretical predictions for ecological speciation where divergence in the face of homogenizing gene flow is promoted by linkage between allelic combinations at loci under selection (clustering, coupling), particularly in genomic areas of low recombination (Butlin & Smadja, 2018; Ortiz-Barrientos & James, 2017; Ravinet et al., 2017; Seehausen et al., 2014; Yeaman et al., 2016). Empirical evidences for these predictions are, thus far, scarce and dominated by systems at early stages of speciation (Feulner et al., 2015; Malinsky et al., 2015; Marques et al., 2017; Meier et al., 2018; Samuk et al., 2017). Later-stage cases appear to be rare, presumably because gene flow is expected to cease rapidly at later stages of speciation leading to rapid genome-wide differentiation (Flaxman et al., 2014; Ravinet et al., 2018; Riesch et al., 2017). The *Silene* system represents such a comparatively advanced stage of speciation with still highly heterogeneous differentiation. The data presented here suggest that this heterogeneity in differentiation is generated by divergent selection driving the evolution of barrier loci.

## Acknowledgements

We are thankful to Emelie Hallander, Karin Steffen and Rasmus Jansson for assistance with plant cultivation and molecular lab work, to Alex Buerkle, Pär Ingvarsson and Martin Lascoux for insightful comments on this work. We are grateful for support from the Science for Life Laboratory and the National Genomics Infrastructure (NGI), Sweden. Computations were performed on resources provided by SNIC through the Uppsala Multidisciplinary Center for Advanced Computational Science (UPPMAX) under projects SNIC 2017/7-406 and uppstore2017241. This research was funded by project grant no. 2012-03622 of the Swedish Research Council (vetenskapsrådet) to SK.

## Author contributions

SK, XL and SG designed the study, XL performed the lab work with the help of lab assistants, XL conducted analyses with input from SK and SG, SK and XL produced figures and wrote the manuscript with contributions from SG.

## Data Accessibility Statement

Double-digest RAD (ddRAD) sequencing data will be available on NCBI’s Short Read Archive (SRA, project number to be added after upload following acceptance). The variant call format (VCF) file and Perl codes for genetic analysis, R codes for plotting and Python codes for demographical modeling will be submitted to dryad (project number to be added after upload following acceptance).

## References

Alexander, D. H., Novembre, J., & Lange, K. (2009). Fast model-based estimation of ancestry in unrelated individuals. Genome Research, 19, 1655–1664.

Berner, D., & Roesti, M. (2017). Genomics of adaptive divergence with chromosome-scale heterogeneity in crossover rate. Molecular Ecology, 26, 6351–6369.

Bolger, A. M., Lohse, M., & Usadel, B. (2014). Trimmomatic: a flexible trimmer for Illumina sequence data. Bioinformatics, 30, 2114–2120.

Burri, R. (2017). Interpreting differentiation landscapes in the light of long-term linked selection. Evolution Letters, 1, 118–131.

Burri, R., Nater, A., Kawakami, T., Mugal, C. F., Olason, P. I., Smeds, L., … Ellegren, H. (2015). Linked selection and recombination rate variation drive the evolution of the genomic landscape of differentiation across the speciation continuum of *Ficedula* flycatchers. Genome Research, 25, 1656–1665.

Butlin, R. K., & Smadja, C. M. (2018). Coupling, reinforcement, and speciation. The American Naturalist, 191, 155–172.

Catchen, J., Hohenlohe, P. A., Bassham, S., Amores, A., & Cresko, W. A. (2013). Stacks: an analysis tool set for population genomics. Molecular Ecology, 22, 3124–3140.

Charlesworth, B. (2009). Fundamental concepts in genetics: effective population size and patterns of molecular evolution and variation. Nature Reviews Genetics, 10, 195–205.

Chong, Z., Ruan, J., & Wu, C.-I. (2012). Rainbow: an integrated tool for efficient clustering and assembling RAD-seq reads. Bioinformatics, 28, 2732–2737.

Christe, C., Stölting, K. N., Paris, M., Fraїsse, C., Bierne, N., & Lexer, C. (2017). Adaptive evolution and segregating load contribute to the genomic landscape of divergence in two tree species connected by episodic gene flow. Molecular Ecology, 26, 59–76.

Cleveland, W. S. (1979). Robust locally weighted regression and smoothing scatterplots. Journal of the American Statistical Association, 74, 829–836.

Coyne, J. A., & Orr, H. A. (2004). Speciation). Sunderland, Massachusetts: Sinauer. 545 pp.

Cruickshank, T. E., & Hahn, M. W. (2014). Reanalysis suggests that genomic islands of speciation are due to reduced diversity, not reduced gene flow. Molecular Ecology, 23, 3133–3157.

Danecek, P., Auton, A., Abecasis, G., Albers, C. A., Banks, E., DePristo, M. A., … Genomes Project Analysis, G. (2011). The variant call format and VCFtools. Bioinformatics, 27, 2156–2158.

Delmore, K. E., Lugo Ramos, J. S., Van Doren, B. M., Lundberg, M., Bensch, S., Irwin, D. E., & Liedvogel, M. (2018). Comparative analysis examining patterns of genomic differentiation across multiple episodes of population divergence in birds. Evolution Letters, 2, 76–87.

Duranton, M., Allal, F., Fraïsse, C., Bierne, N., Bonhomme, F., & Gagnaire, P.-A. (2018). The origin and remolding of genomic islands of differentiation in the European sea bass. Nature Communications, 9, 2518.

Favre, A., Widmer, A., & Karrenberg, S. (2017). Differential adaptation drives ecological speciation in campions (*Silene*): evidence from a multi-site transplant experiment. New Phytologist, 213, 1487–1499.

Feulner, P. G. D., Chain, F. J. J., Panchal, M., Huang, Y., Eizaguirre, C., Kalbe, M., … Milinski, M. (2015). Genomics of divergence along a continuum of parapatric population differentiation. PLoS Genetics, 11, e1004966.

Flaxman, S. M., Wacholder, A. C., Feder, J. L., & Nosil, P. (2014). Theoretical models of the influence of genomic architecture on the dynamics of speciation. Molecular Ecology, 23, 4074–4088.

Friedrich, H. C. (1979). Caryophyllaceae. In K. H. Rechinger (Ed.), Illustrierte Flora von Mitteleuropa (2 ed.). Hamburg: Parey.

Garrison, E. (2012). Vcflib: A C++ library for parsing and manipulating VCF files. Retrieved from: https://github.com/ekg/vcflib

Garrison, E., & Marth, G. (2012). Haplotype-based variant detection from short-read sequencing. arXiv preprint, arXiv:1207.3907.

Goudet, J. (2004). *hierfstat*, a package for R to compute and test hierarchical F–statistics. Molecular Ecology Notes, 5, 184–186.

Goulson, D., & Jerrim, K. (1997). Maintenance of the species boundary between *Silene dioica* and *S. latifolia* (red and white campion). Oikos, 79, 115–126.

Guirao-Rico, S., Sánchez-Gracia, A., & Charlesworth, D. (2017). Sequence diversity patterns suggesting balancing selection in partially sex-linked genes of the plant *Silene latifolia* are not generated by demographic history or gene flow. Molecular Ecology, 26, 1357–1370.

Gutenkunst, R. N., Hernandez, R. D., Williamson, S. H., & Bustamante, C. D. (2009). Inferring the joint demographic history of multiple populations from multidimensional SNP frequency data. PLoS Genetics, 5, e1000695.

Han, F., Lamichhaney, S., Grant, B. R., Grant, P. R., Andersson, L., & Webster, M. T. (2017). Gene flow, ancient polymorphism, and ecological adaptation shape the genomic landscape of divergence among Darwin’s finches. Genome Research, 27, 1004–1015.

Hathaway, L., Malm, J. U., & Prentice, H. C. (2009). Geographically congruent large-scale patterns of plastid haplotype variation in the European herbs *Silene dioica* and *S. latifolia* (Caryophyllaceae). Botanical Journal of the Linnean Society, 161, 153–170.

Hewitt, G. (2000). The genetic legacy of the Quaternary ice ages. Nature, 405, 907–913.

Hothorn, T., Hornik, K., Wiel, M. A. v. d., & Zeileis, A. (2006). A Lego system for conditional inference. The American Statistician, 60, 257–263.

Hu, X. S., & Filatov, D. A. (2015). The large-X effect in plants: Increased species divergence and reduced gene flow on the *Silene* X-chromosome. Molecular Ecology, 25, 2609–2619.

Karrenberg, S., & Favre, A. (2008). Genetic and ecological differentiation in the hybridizing campions *Silene dioica* and *S. latifolia*. Evolution, 62, 763–773.

Karrenberg, S., Liu, X., Hallander, E., Favre, A., Herforth-Rahmé, J., & Widmer, A. (2019). Ecological divergence plays an important role in strong but complex reproductive isolation in campions (*Silene*). Evolution, 73, 245–261.

Krasovec, M., Chester, M., Ridout, K., & Filatov, D. A. (2018). The mutation rate and the age of the sex chromosomes in *Silene latifolia*. Current Biology, 28, 1832–1838.

Lengerova, M., Kejnovsky, E., Hobza, R., Macas, J., Grant, S. R., & Vyskot, B. (2004). Multicolor FISH mapping of the dioecious model plant, *Silene latifolia*. Theoretical and Applied Genetics, 108, 1193–1199.

Li, H. (2013). Aligning sequence reads, clone sequences and assembly contigs with BWA-MEM. arXiv preprint, arXiv:1303.3997.

Liu, X., & Karrenberg, S. (2018). Genetic architecture of traits associated with reproductive barriers in *Silene*: Coupling, sex chromosomes and variation. Molecular Ecology, 27, 3889–3904.

Malinsky, M., Challis, R. J., Tyers, A. M., Schiffels, S., Terai, Y., Ngatunga, B. P., … Turner, G. F. (2015). Genomic islands of speciation separate cichlid ecomorphs in an East African crater lake. Science, 350, 1493–1498.

Marques, D. A., Lucek, K., Haesler, M. P., Feller, A. F., Meier, J. I., Wagner, C. E., … Seehausen, O. (2017). Genomic landscape of early ecological speciation initiated by selection on nuptial colour. Molecular Ecology, 26, 7–24.

Martin, S. H., Dasmahapatra, K. K., Nadeau, N. J., Salazar, C., Walters, J. R., Simpson, F., … Jiggins, C. D. (2013). Genome-wide evidence for speciation with gene flow in *Heliconius* butterflies. Genome Research, 23, 1817–1828.

Meier, J. I., Marques, D. A., Wagner, C. E., Excoffier, L., & Seehausen, O. (2018). Genomics of parallel ecological speciation in Lake Victoria cichlids. Molecular Biology and Evolution, 35, 1489–1506.

Minder, A. M., Rothenbuehler, C., & Widmer, A. (2007). Genetic structure of hybrid zones between *Silene latifolia* and *Silene dioica* (Caryophyllaceae): evidence for introgressive hybridization. Molecular Ecology, 16, 2504–2516.

Muir, G., Dixon, C. J., Harper, A. L., & Filatov, D. A. (2012). Dynamics of drift, gene flow, and selection during speciation in *Silene*. Evolution, 66, 1447–1458.

Nachman, M. W., & Payseur, B. A. (2012). Recombination rate variation and speciation: theoretical predictions and empirical results from rabbits and mice. Philosophical Transactions of the Royal Society B: Biological Sciences, 367, 409–421.

Nielsen, R. (2005). Molecular signatures of natural selection. Annual Review of Genetics, 39, 197–218.

O’Leary, S. J., Puritz, J. B., Willis, S. C., Hollenbeck, C. M., & Portnoy, D. S. (2018). These aren’t the loci you’e looking for: Principles of effective SNP filtering for molecular ecologists. Molecular Ecology, 27, 3193–3206.

Oksanen, J., Blanchet, F. G., Friendly, M., Kindt, R., Legendre, P., McGlinn, D., … Wagner, H. (2019). vegan: Community Ecology Package. R package version 2.5–5. Retrieved from: https://CRAN.R-project.org/package=vegan

Ortiz-Barrientos, D., & James, M. E. (2017). Evolution of recombination rates and the genomic landscape of speciation. Journal of Evolutionary Biology, 30, 1519–1521.

Pease, J. B., Haak, D. C., Hahn, M. W., & Moyle, L. C. (2016). Phylogenomics reveals three sources of adaptive variation during a rapid radiation. PLoS Biology, 14, e1002379.

Peterson, B. K., Weber, J. N., Kay, E. H., Fisher, H. S., & Hoekstra, H. E. (2012). Double digest RADseq: an inexpensive method for de novo SNP discovery and genotyping in model and non-model species. PLoS ONE, 7, e37135.

Poelstra, J. W., Vijay, N., Bossu, C. M., Lantz, H., Ryll, B., Müller, I., … Wolf, J. B. W. (2014). The genomic landscape underlying phenotypic integrity in the face of gene flow in crows. Science, 344, 1410–1414.

Prentice, H. C., Malm, J. U., & Hathaway, L. (2008). Chloroplast DNA variation in the European herb *Silene dioica* (red campion): postglacial migration and interspecific introgression. Plant Systematics and Evolution, 272, 23–37.

Purcell, S., Neale, B., Todd-Brown, K., Thomas, L., Ferreira, M. A. R., Bender, D., … Sham, P. C. (2007). PLINK: a tool set for whole-genome association and population-based linkage analyses. American Journal of Human Genetics, 81, 559–575.

Puritz, J. B., Hollenbeck, C. M., & Gold, J. R. (2014). dDocent: a RADseq, variant-calling pipeline designed for population genomics of non-model organisms. PeerJ, 2, e431.

Rafajlović, M., Emanuelsson, A., Johannesson, K., Butlin, R. K., & Mehlig, B. (2016). A universal mechanism generating clusters of differentiated loci during divergence-with-migration. Evolution, 70, 1609–1621.

Rautenberg, A., Hathaway, L., Oxelman, B., & Prentice, H. C. (2010). Geographic and phylogenetic patterns in *Silene* section *Melandrium* (Caryophyllaceae) as inferred from chloroplast and nuclear DNA sequences. Molecular Phylogenetics and Evolution, 57, 978–991.

Ravinet, M., Faria, R., Butlin, R. K., Galindo, J., Bierne, N., Rafajlović, M., … Westram, A. M. (2017). Interpreting the genomic landscape of speciation: a road map for finding barriers to gene flow. Journal of Evolutionary Biology, 30, 1450–1477.

Ravinet, M., Yoshida, K., Shigenobu, S., Toyoda, A., Fujiyama, A., & Kitano, J. (2018). The genomic landscape at a late stage of stickleback speciation: High genomic divergence interspersed by small localized regions of introgression. PLOS Genetics, 14, e1007358.

Renaut, S., Grassa, C. J., Yeaman, S., Moyers, B. T., Lai, Z., Kane, N. C., … Rieseberg, L. H. (2013). Genomic islands of divergence are not affected by geography of speciation in sunflowers. Nature Communications, 4, 1827.

Richards, E. J., Servedio, M. R., & Martin, C. H. (2019). Searching for sympatric speciation in the genomic era. Bioessays, 41, e1900047.

Riesch, R., Muschick, M., Lindtke, D., Villoutreix, R., Comeault, A. A., Farkas, T. E., … Nosil, P. (2017). Transitions between phases of genomic differentiation during stick-insect speciation. Nat Ecol Evol, 1, 82.

Rockman, M. V. (2012). The QTN program and the alleles that matter for evolution: All that’s gold does not glitter. Evolution, 66, 1–17.

Rougemont, Q., & Bernatchez, L. (2018). The demographic history of Atlantic salmon (*Salmo salar*) across its distribution range reconstructed from approximate Bayesian computations*. Evolution, 72, 1261–1277.

Roux, C., Fraïsse, C., Castric, V., Vekemans, X., Pogson, G. H., & Bierne, N. (2014). Can we continue to neglect genomic variation in introgression rates when inferring the history of speciation? A case study in a *Mytilus* hybrid zone. Journal of Evolutionary Biology, 27, 1662–1675.

Roux, C., Fraisse, C., Romiguier, J., Anciaux, Y., Galtier, N., & Bierne, N. (2016). Shedding light on the grey zone of speciation along a continuum of genomic divergence. PLoS Biology, 14, e2000234.

Sambatti, J. B., Strasburg, J. L., Ortiz-Barrientos, D., Baack, E. J., & Rieseberg, L. H. (2012). Reconciling extremely strong barriers with high levels of gene exchange in annual sunflowers. Evolution, 66, 1459–1473.

Samuk, K., Owens, G. L., Delmore, K. E., Miller, S. E., Rennison, D. J., & Schluter, D. (2017). Gene flow and selection interact to promote adaptive divergence in regions of low recombination. Molecular Ecology, 26, 4378–4390.

Seehausen, O., Butlin, R. K., Keller, I., Wagner, C. E., Boughman, J. W., Hohenlohe, P. A., … Widmer, A. (2014). Genomics and the origin of species. Nature Reviews: Genetics, 15, 176–192.

Tine, M., Kuhl, H., Gagnaire, P.-A., Louro, B., Desmarais, E., Martins, R. S. T., … Reinhardt, R. (2014). European sea bass genome and its variation provide insights into adaptation to euryhalinity and speciation. Nature Communications, 5, 5770.

Vijay, N., Weissensteiner, M., Burri, R., Kawakami, T., Ellegren, H., & Wolf, J. B. W. (2017). Genomewide patterns of variation in genetic diversity are shared among populations, species and higher-order taxa. Molecular Ecology, 26, 4284–4295.

Wang, J., Street, N. R., Scofield, D. G., & Ingvarsson, P. K. (2016). Variation in linked selection and recombination drive genomic divergence during allopatric speciation of european and american aspens. Molecular Biology and Evolution, 33, 1754–1767.

Wolf, J. B. W., & Ellegren, H. (2017). Making sense of genomic islands of differentiation in light of speciation. Nature Reviews: Genetics, 18, 87–100.

Yeaman, S. (2015). Local adaptation by alleles of small effect. The American Naturalist, 186, S74–S89.

Yeaman, S., Aeschbacher, S., & Bürger, R. (2016). The evolution of genomic islands by increased establishment probability of linked alleles. Molecular Ecology, 25, 2542–2558.

