## Supporting Information for "Evolution of barrier loci at an intermediate stage of speciation with gene flow"

### Supplementary Figures

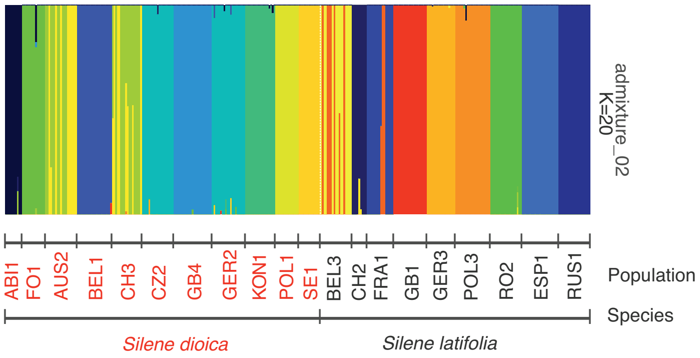


**Figure S1.** Barplot of a structure analysis of the closely related campions *Silene dioica* (11 populations) and *S. latifolia* (9 populations) using $K = 20$, the total number of populations.

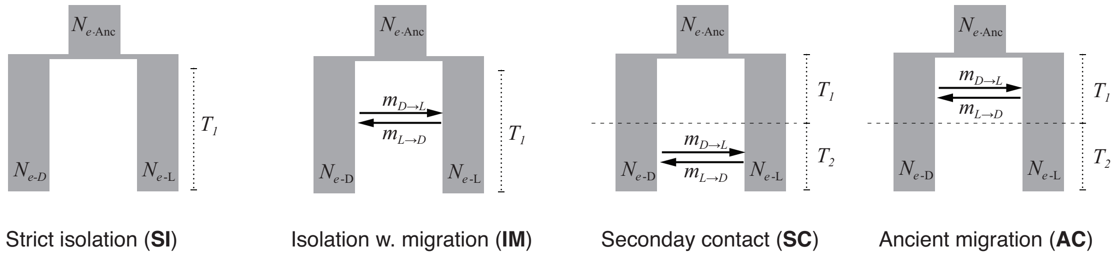


**Figure S2.** Illustration of four basic demographic scenarios: strict isolation (SI); isolation with migration (IM), in which divergence evolves with continuous gene flow; ancient migration (AM), in which divergence occurs initially with gene flow and later gene flow ceases; secondary contact (SC), in which divergence occurs initially without gene flow and later gene flow occurs. D and L subscripts stand for the two species modeled here, the campions, *Silene dioica* (D) and *S. latifolia* (L).

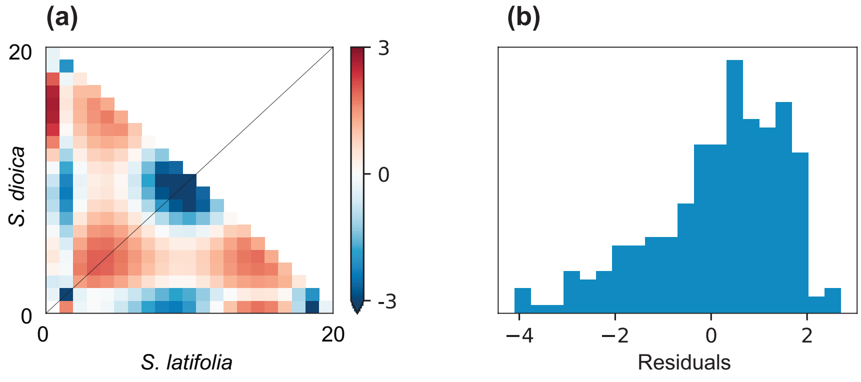


**Figure S3.** Residuals of the best-supported demographic model of the lineage divergence between the campions *Silene dioica* and *S. latifolia*, isolation with migration including heterogeneous migration rate and population size (IMhmhn); **(a)** heat map and **(b)** frequency distribution of deviations in SNP numbers between the observed joint site frequency spectrum (SFS) and the SFS generated under the best-supported model.

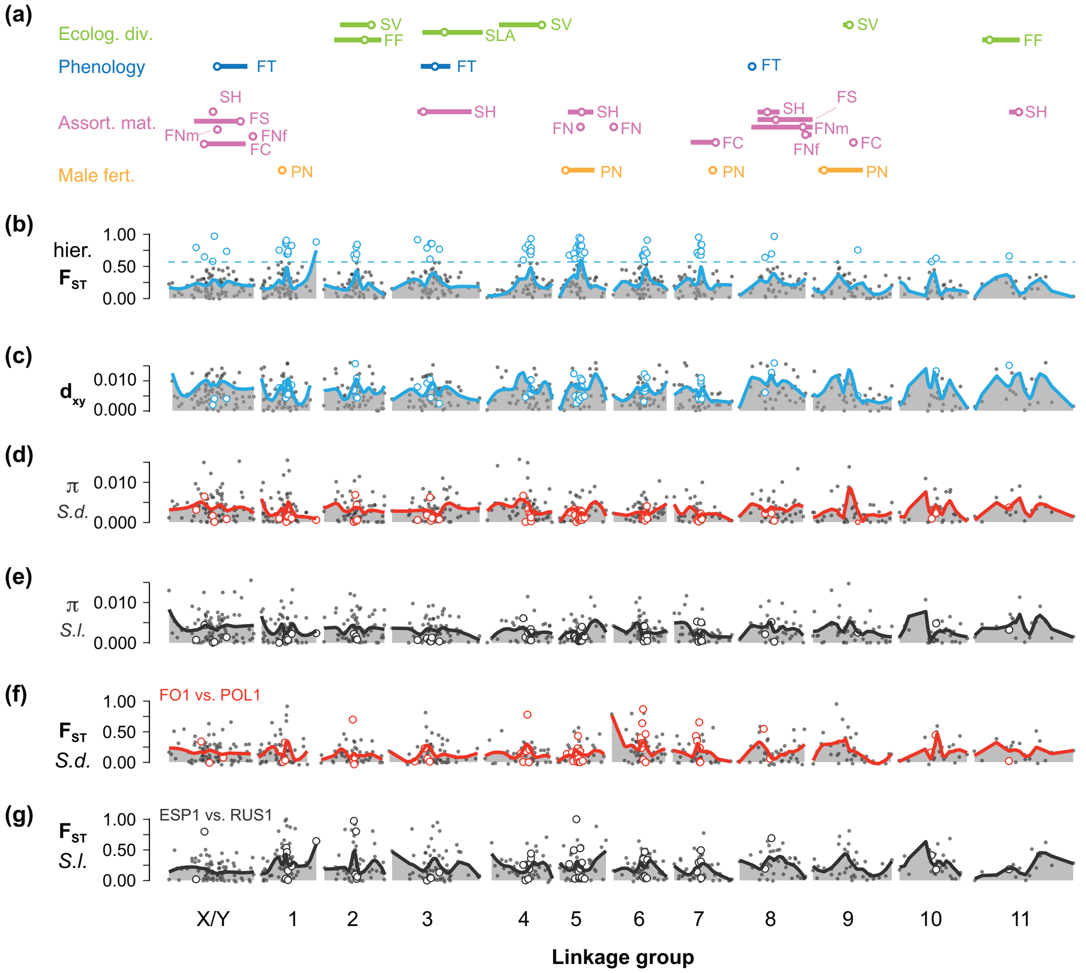


**Figure S4.** Genomic landscapes and QTLs for traits associated with reproductive barriers between the campions *Silene dioica* and *S. latifolia* based on a linkage map of the $F_{2LD}$ cross (for the $F_{2DL}$ cross see main Figure 4); **(a)**, locations (circles) and 1.5 LOD drop intervals (lines) of QTLs related ecological divergence (FF: first-year flowering, SUC: leaf succulence, SLA : specific leaf area), phenology (FT: flowering time) and assortative mating (FC: flower color, FS: flower size, FNf and FNm: flower number for females and for males, SH: stem height), hybrid male fertility reduction (PN: pollen number); **(b)**, hierarchical $F_{ST}$ between the two species with the 95% quantile of overall $F_{ST}$ distribution (dashed line); **(c)**, sequence divergence ($d_{XY}$); **(d)** and **(e)**, nucleotide diversity (π) within *S. dioica* (*S.d.*) and within *S. latifolia* (*S.l.*); **(f)** and **(g)**, $F_{ST}$ of the most differentiated population pair within *S. dioica* and within *S. latifolia*. Lines in panels **(b)** - **(g)** are drawn using LOESS functions; empty circles represent contigs with hierarchical $F_{ST}$ exceeding the 95% quantile (genomic islands of differentiation).

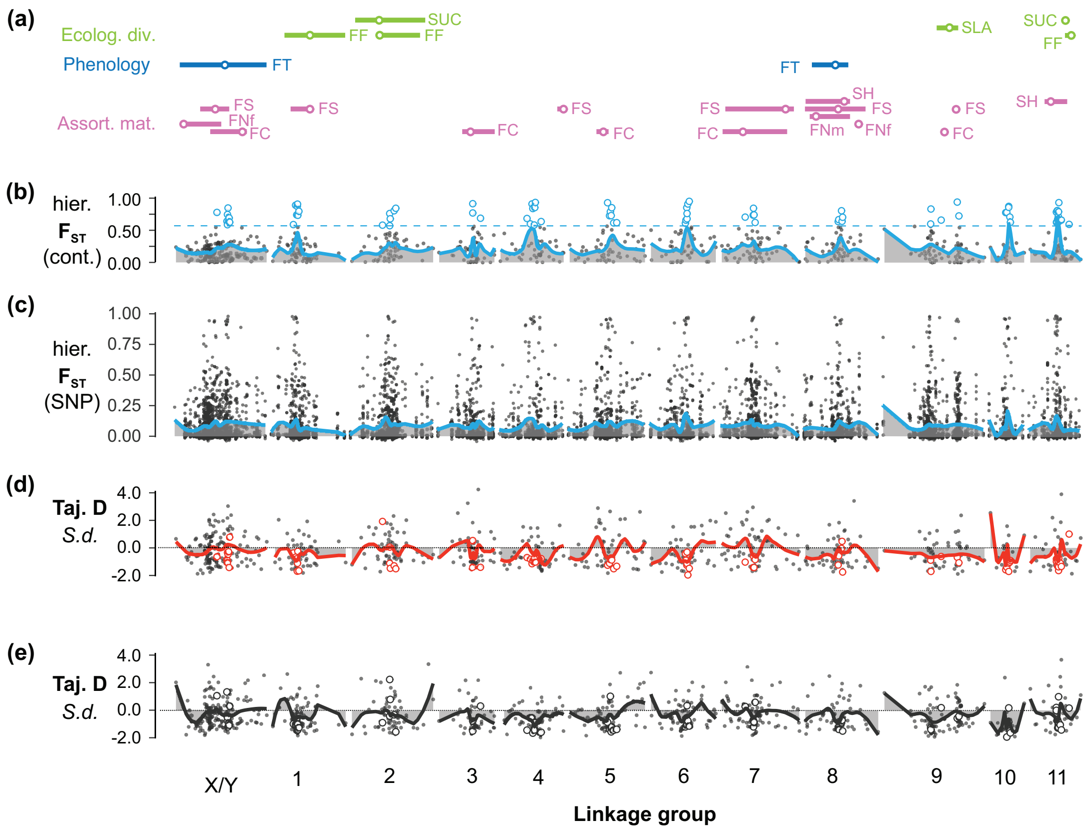


**Figure S5.** Genomic landscapes and QTLs for traits associated with reproductive barriers between the campions *Silene dioica* and *S. latifolia* based on a linkage map of the $F_{2DL}$ cross; **(a)**, locations (circles) and 1.5 LOD drop intervals (lines) of QTLs related ecological divergence (FF: first-year flowering, SUC: leaf succulence, SLA: specific leaf area), phenology (FT: flowering time) and assortative mating (FC: flower color, FS: flower size, FNf and FNm: flower number for females and for males, SH: stem height); **(b)**, contig-based hierarchical $F_{ST}$ between the two species with the 95% quantile of overall $F_{ST}$ distribution (dashed line); **(c)** SNP-based hierarchical $F_{ST}$ between the two species. **(d)** and **(e)** Tajima's D within *S. dioica* (*S.d.*) and within *S. latifolia* (*S.l.*). Lines in the panels **(b)** - **(e)** are drawn using LOESS functions and empty circles represent contigs with hierarchical $F_{ST}$ exceeding the 95% quantile (genomic islands of differentiation).

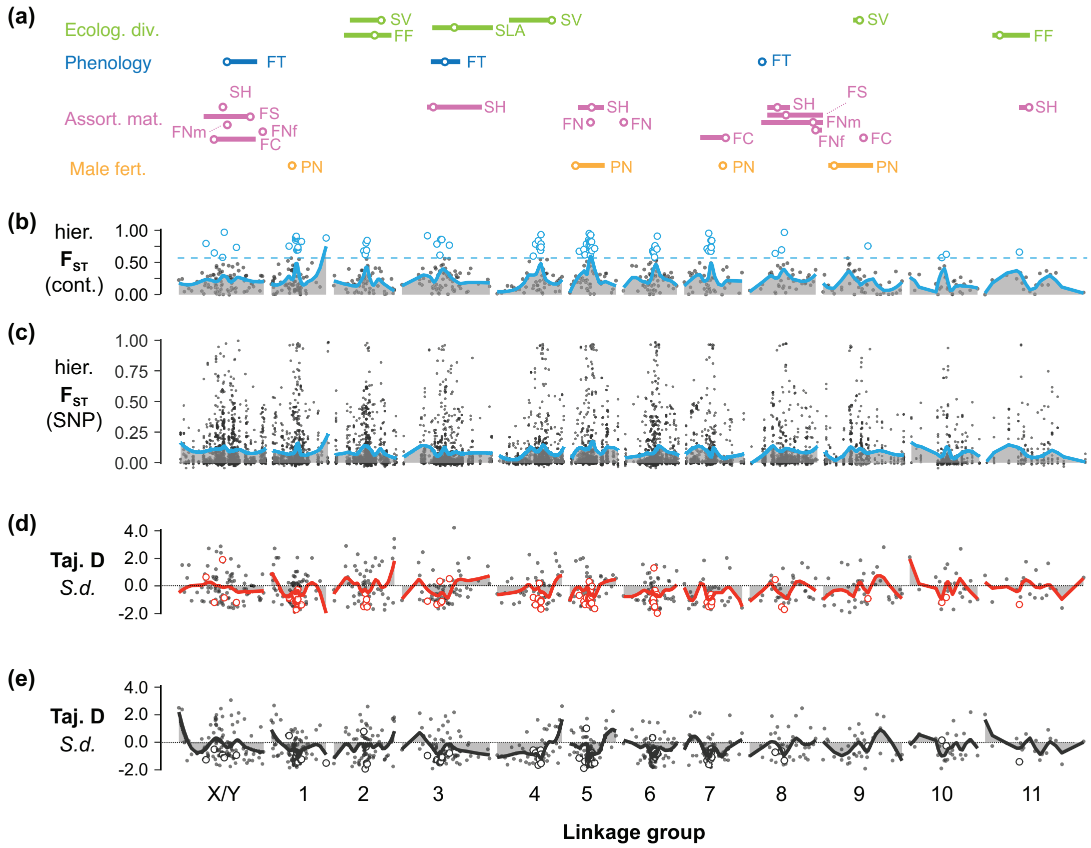


**Figure S6.** Genomic landscapes and QTLs for traits associated with reproductive barriers between the campions *Silene dioica* and *S. latifolia* based on a linkage map of the $F_{2LD}$ cross; **(a)**, locations (circles) and 1.5 LOD drop intervals (lines) of QTLs related ecological divergence (FF: first-year flowering, SUC: leaf succulence, SLA: specific leaf area), phenology (FT: flowering time) and assortative mating (FC: flower color, FS: flower size, FNf and FNm: flower number for females and for males, SH: stem height), and hybrid male fertility reduction (PN: pollen number); **(b)**, contig-based hierarchical $F_{ST}$ between the two species with the 95% quantile of overall $F_{ST}$ distribution (dashed line); **(c)** SNP-based hierarchical $F_{ST}$ between the two species. **(d)** and **(e)** Tajima's D within *S. dioica* (*S.d.*) and within *S. latifolia* (*S.l.*). Lines in the panels **(b)** - **(e)** are drawn using LOESS functions and empty circles represent contigs with hierarchical $F_{ST}$ exceeding the 95% quantile (genomic islands of differentiation).

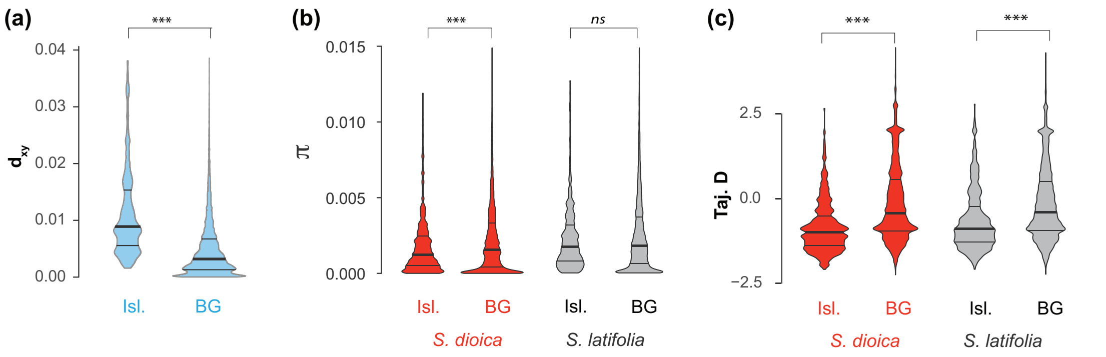


**Figure S7.** Violin plots for population genetic statistics in the campions *Silene dioica* and *S. latifolia* in genomic islands of differentiation (Isl.) and the genomic background (BG) for all contigs in the dataset **(a)** between-species $d_{XY}$. **(b)** genetic diversity (π), and **(c)** for Tajima's D. Within each violin, the horizontal lines represent the 25%, 50% (median, thick lines) and 75% quantiles. Results from Mood's median tests are given on the top (***, P-Value < 0.001, *ns*, P > 0.05).

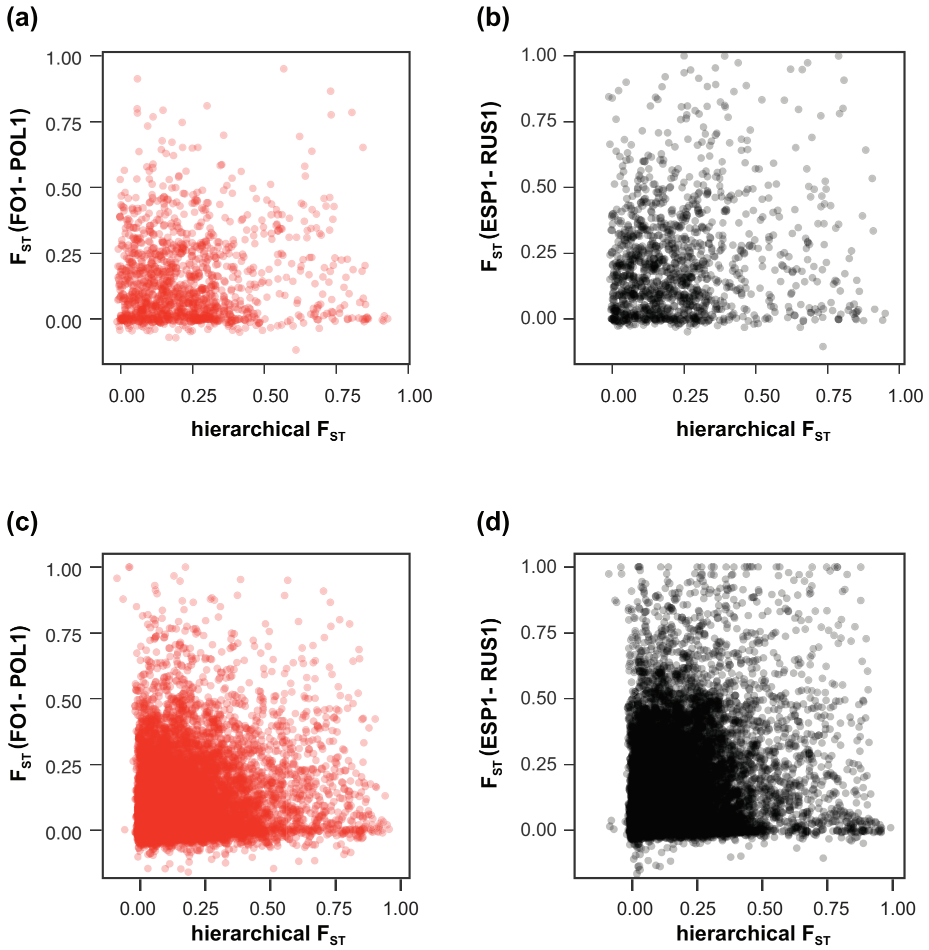


**Figure S8.** Scatterplots of hierarchical $F_{ST}$ between the campions *Silene dioica* and *S. latifolia* (x-axis) against $F_{ST}$ of the most diverged population pair within each species (y-axis), FO1-POL1 for *S. dioica* **(a, c)** and ESP1-RUS1 for *S. latifolia* **(b, c)**. Panels **(a)** and **(b)** represent mapped contigs and panels **(c)** and **(d)** all contigs in the dataset.

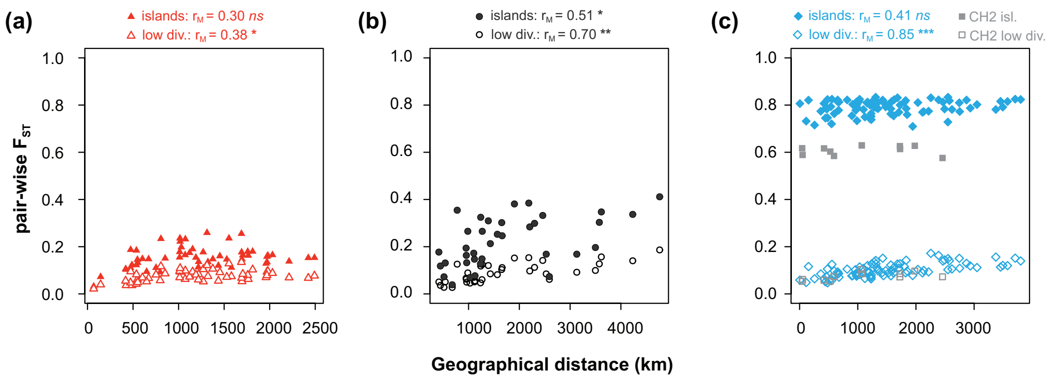


**Figure S9.** Scatterplots of pair-wise differentiation ($F_{ST}$) against geographic distance (km) for population pairs within the campion *Silene dioica* **(a)**, within *S. latifolia* **(b)**, and between the two species **(c)** based on all contigs in the dataset. Pair-wise $F_{ST}$ is given for between-species differentiation islands (filled symbols(> 95% quantile) and for regions of low between-species divergence (empty symbols, < 25% quantile) based on mapped contigs. Results of Mantel tests are indicates at the top; *ns*, $P > 0.05$; *, $P < 0.05$; **, $P < 0.01$; ***, $P < 0.001$.
