## Supporting Information for "Evolution of barrier loci at an intermediate stage of speciation with gene flow"

**Table S1.** Study populations of *Silene dioica* and *S. latifolia*, latitude and longitude are given in decimal degrees (WGS84).

| Population | Species | Location | Latitud e | Longitude | Collector | No. of faimilies | No. of individuals |
|---|---|---|---|---|---|---|---|
| ABI1 | *Silene dioica* | Sweden | 68.35 | 18.820 | Tora Finderup Nilsen | 10 | 10 |
| AUS2 | *Silene dioica* | Austria | 47.064 | 9.850 | Mark van Kleunen | 11 | 19 |
| BEL1 | *Silene dioica* | Belgium | 49.593 | 5.576 | Fabienne van Rossum | 12 | 21 |
| BEL3 | *Silene latifolia* | Belgium | 49.548 | 5.594 | | 12 | 20 |
| CH2 | *Silene latifolia* | Switzerland | 46.814 | 9.400 | Christian Rixen | 9 | 9 |
| CH3 | *Silene dioica* | Switzerland | 46.450 | 9.830 | | 11 | 18 |
| CZ2 | *Silene dioica* | Czech Repulic | 50.768 | 14.790 | Michael Sochor | 8 | 19 |
| ESP1 | *Silene latifolia* | Spain | 40.626 | -3.894 | Luis Giminez | 11 | 22 |
| FO1 | *Silene dioica* | Faroe island | 62.026 | -6.772 | Anna Maria Fosaa | 3 | 13 |
| FRA1 | *Silene latifolia* | France | 47.586 | -3.110 | Fabienne van Rossum | 10 | 16 |
| GB1 | *Silene latifolia* | Great Britain | 57.687 | -1.840 | Deborah Charlesworth | 12 | 20 |
| GB4 | *Silene dioica* | Great Britain | 53.384 | -1.538 | | 12 | 23 |
| GER2 | *Silene dioica* | Germany | 51.006 | 12.773 | Adrien Favre | 11 | 20 |
| GER3 | *Silene latifolia* | Germany | 51.297 | 11.224 | | 10 | 17 |
| KON1 | *Silene dioica* | Norway | 62.303 | 9.605 | Tora Finderup Nilsen | 10 | 18 |

| | | | | | | | |
|---|---|---|---|---|---|---|---|
| POL1 | *Silene dioica* | Poland | 54.412 | 18.415 | Maria Uscka-Perzanowska | 8 | 14 |
| POL3 | *Silene latifolia* | Poland | 53.041 | 18.604 | | 11 | 21 |
| RO2 | *Silene latifolia* | Romania | 45.738 | 25.750 | Andreea Spinu | 8 | 19 |
| RUS1 | *Silene latifolia* | Russia | 55.186 | 58.614 | Vladimir Semerikov/ Natalia Kutlunina | 19 | 19 |
| SE1 | *Silene dioica* | Sweden | 61.627 | 17.446 | Adrian Pop | 8 | 13 |

**Table S2.** Comparison of demographic models of the lineage split between the campions *Silene dioica* and *S. latifolia.* Basic models are strict isolation (SI), isolation with migration (IM), secondary contact (SC) and ancient migration (AM). Further parameter were added to these model to allow for population sizechange (exp.), heterogeneous migration rates (hm), heterogeneous effective population size (hn) and for heterogeneiety in both migration rate and population size (hmhn). Given are the number of parameters (K), the likelihood (L) , Akaikes information criterion (AIC) and P-values for likelihood ratio tests within nested models.

| **Models** | **K** | **L** | **AIC** | ***P*-value** | | |
|---|---|---|---|---|---|---|
| | | | | **vs. basic** | **vs. hm** | **vs. hn** |
| SI_basic | 3 | -837.77 | 1681.55 | | | |
| SI_exp | 5 | -816.54 | 1643.07 | $5.98\times10^{-10}$ | | |
| SI_hn | 5 | -723.24 | 1456.48 | 0 | | |
| IM_basic | 4 | -761.45 | 1530.90 | | | |
| IM_exp | 6 | -752.73 | 1517.47 | $1.64\times10^{-4}$ | | |
| IM_hm | 5 | -700.78 | 1411.57 | 0 | | |
| IM_hn | 6 | -696.21 | 1404.42 | 0 | | |
| IM_hmhn | 8 | -685.72 | 1387.45 | 0 | $1.30\times10^{-6}$ | $2.79\times10^{-5}$ |
| SC_basic | 5 | -761.45 | 1532.89 | | | |
| SC_exp | 7 | -752.15 | 1518.30 | $9.19\times10^{-5}$ | | |
| SC_hm | 6 | -700.81 | 1413.63 | 0 | | |
| SC_hn | 7 | -696.67 | 1407.34 | 0 | | |
| SC_hmhn | 9 | -685.73 | 1389.45 | 0 | $1.27\times10^{-6}$ | $1.77\times10^{-5}$ |
| AM_basic | 5 | -761.52 | 1533.04 | | | |
| AM_exp | 7 | -752.76 | 1519.53 | $1.58\times10^{-4}$ | | |
| AM_hm | 6 | -700.78 | 1413.57 | 0 | | |
| AM_hn | 7 | -704.00 | 1422.01 | 0 | | |
| AM_hmhn | 9 | -685.78 | 1389.56 | 0 | $1.37\times10^{-6}$ | $1.22\times10^{-8}$ |

**Table S3.** Summary of parameter estimates for demographic models of the lineage split between the campions *Silene dioica* and *S. latifolia* in ∂a∂i. Four basic scenarios, strict isolation (SI), isolation with migration (IM), secondary contact (SC) and ancient migration (AM) were modeled ("basic"). To these models, we added population expansion ("exp"), heterogeneous migration rates ("hm"), heterogeneous population size ("hn") and both heterogeneous migration rate and heterogeneous population size("hmhn"). Point estimates of theta, population sizes, divergence times (T) in units of 2 x $N_{e\text{-}Anc}$, overall population migration rates (M) in units of 2 × $N_{e\text{-}Anc}$ × m, population size change factors (b), population size is reduction factor ($N_{e1}$ / $N_{e2}$) where two population sizes are modeled, and percentages of different loci. Subscripts refer to estimates for the two species, *S. dioica* (D) and *S. latifolia* (L).

| Models | Theta | $N_{e\text{-}Anc}$ | $N_{e\text{-}D}$ | $N_{e\text{-}L}$ | Divergence time | | M | $b_D$ | $b_L$ | $N_{e1}$ / $N_{e2}$ | Percentage of loci | | | |
|---|---|---|---|---|---|---|---|---|---|---|---|---|---|---|
| | | | | | | | | | | | $M = 0$ | | $M > 0$ | |
| | | | | | $T_1$ * | $T_2$ ** | | | | | $N_{e1}$ | $N_{e2}$ | $N_{e1}$ | $N_{e2}$ |
| SI_basic | 1298.6 | 401211.3 | 0.588 | 0.801 | 0.205 | | | | | | | | | |
| SI_exp | 1366.7 | 422246.0 | 0.190 | 0.391 | 0.157 | | | 6.104 | 2.790 | | | | | |
| SI_hn | 1319.9 | 407786.7 | 3.429 | 4.999 | 0.187 | | | | | 9.11 | 20.91% | 79.09% | | |
| IM_basic | 1202.4 | 371495.1 | 0.647 | 0.875 | 0.338 | | 0.430 | | | | | | | |
| IM_exp | 1273.6 | 393484.4 | 0.297 | 0.573 | 0.262 | | 0.391 | 3.010 | 1.704 | | | | | |
| IM_hm | 1170.0 | 361485.9 | 0.669 | 0.912 | 0.383 | | 2.442 | | | | 59.56% | | 40.44% | |
| IM_hn | 1274.7 | 393813.6 | 2.014 | 2.866 | 0.257 | | 0.392 | | | 5.29 | | | 25.99% | 74.01% |
| IM_hmhn | 1225.3 | 378576.0 | 2.713 | 3.791 | 0.302 | | 1.379 | | | 5.42 | 0.04% | 52.04% | 13.56% | 34.36% |
| SC_basic | 1202.3 | 371471.0 | 0.647 | 0.875 | 0.005 | 0.334 | 0.431 | | | | | | | |
| SC_exp | 1272.1 | 393021.8 | 0.375 | 0.643 | 0.071 | 0.190 | 0.394 | 2.449 | 1.542 | | | | | |
| SC_hm | 1167.9 | 360829.3 | 0.671 | 0.914 | 0.023 | 0.362 | 2.416 | | | | 59.26% | | 40.74% | |
| SC_hn | 1279.1 | 395197.6 | 1.664 | 2.369 | 0.000 | 0.257 | 0.397 | | | 4.54 | | | 30.03% | 69.97% |
| SC_hmhn | 1226.4 | 378898.3 | 2.640 | 3.690 | 0.012 | 0.289 | 1.346 | | | 5.31 | 0.00% | 51.68% | 14.07% | 34.25% |

| | | | | | | | | | | | | | | |
|---|---|---|---|---|---|---|---|---|---|---|---|---|---|---|
| AM_basic | 1192.4 | 368391.9 | 0.647 | 0.875 | 0.339 | 0.000 | 0.433 | | | | | | | |
| AM_exp | 1275.4 | 394030.4 | 0.292 | 0.563 | 0.261 | 0.000 | 0.391 | 3.091 | 1.747 | | | | | |
| AM_hm | 1169.1 | 361189.7 | 0.670 | 0.913 | 0.383 | 0.000 | 2.426 | | | | 59.38% | | 40.62% | |
| AM_hn | 1277.6 | 394709.9 | 2.967 | 2.967 | 0.246 | 0.011 | 0.435 | | | 7.64 | | | 19.82% | 80.18% |
| AM_hmhn | 1227.1 | 379122.5 | 2.563 | 3.593 | 2.993 | 0.000 | 1.304 | | | 5.20 | 0.00% | 51.14% | 14.64% | 34.22% |

* For the IM and SI models, $T_1$ is the total divergence time. For the SC models, T1 refers to the time of lineage-split without migration while for the AM models, T1 refers to the time of lineage-split with migration.

** For the SC models, $T_2$ refers to the time of lineage-split with migration while for the AM models, $T_2$ refers to the time of lineage-split without migration.

**Table S4.** Median sequence divergence ($d_{XY}$) between *Silene dioica* and *S. latifolia*, and of nucleotide diversity (π) and Tajima's D within each species for genomic islands of between spcies differentiation and the remaining loci (Background) with Z and *P* values from Mood's median test are given for contigs mapped to linkage maps (a) and for all contigs in the dataset (b) .P-Values significant at alpha = 0.05 are in bold type.

**(a) Mapped contigs**

| | Median | | *Z* | P-value |
|---|---|---|---|---|
| | Islands | Background | | |
| $d_{XY}$ between species | 0.0082 | 0.0049 | 5.80 | **$6.44\times10^{-9}$** |
| π *S. dioica* | 0.0011 | 0.0022 | -4.62 | **$3.85\times10^{-6}$** |
| π *S. latifolia* | 0.0014 | 0.0024 | -4.07 | **$4.80\times10^{-5}$** |
| Tajima's D *S. dioica* | -1.055 | -0.460 | -7.82 | **$5.16\times10^{-15}$** |
| Tajima's D *S. latifolia* | -1.045 | -0.556 | -5.79 | **$6.88\times10^{-9}$** |

**(b) All contigs**

| | Median | | *Z* | P-value |
|---|---|---|---|---|
| | Islands | Background | | |
| $d_{XY}$ between species | 0.0088 | 0.0032 | 19.75 | **$< 2.2\times10^{-16}$** |
| π *S. dioica* | 0.0012 | 0.0014 | -2.84 | **0.0045** |
| π *S. latifolia* | 0.0017 | 0.0017 | -0.46 | 0.6484 |
| Tajima's D *S. dioica* | -0.992 | -0.429 | -11.97 | **$< 2.2\times10^{-16}$** |
| Tajima's D *S. latifolia* | -0.887 | -0.404 | -9.80 | **$< 2.2\times10^{-16}$** |

**Table S5.** Results of Mantel's tests between genetic distance (pairwise $F_{ST}$) and geographical distance for population pairs within and between the campions *Silene dioica* and *S. latifolia* for genomic islands of high differentiation and for regions of low differentitation (above the 95% quantile and below 25% quantile of hierarchical $F_{ST}$ between species, respectively). **(a)** for contigs mapped to linkage maps, **(b)**for all contigs in the dataset. P-Values significant at alpha = 0.05 are in bold type.

**(a) Mapped contigs**

| | **within *S. dioica*** | | **within *S. latifolia*** | | **between species** | |
|---|---|---|---|---|---|---|
| | Islands | low differentiation regions | Islands | low differentiation regions | Islands | low differentiation regions |
| **Mantel *r*** | 0.2195 | 0.2576 | 0.5589 | 0.7601 | 0.3215547 | 0.8984 |
| **P-value** | 0.1216 | 0.1042 | **0.0126** | **0.0002** | 0.1549 | **0.0001** |

**(b) All contigs**

| | **within *S. dioica*** | | **within *S. latifolia*** | | **between species** | |
|---|---|---|---|---|---|---|
| | Islands | low differentiation regions | Islands | low differentiation regions | Islands | low differentiation regions |
| **Mantel *r*** | 0.2981 | 0.3836 | 0.5196 | 0.7008 | 0.4166 | 0.8538 |
| **P-value** | 0.0727 | **0.0163** | **0.0484** | **0.0041** | 0.0828 | **0.0001** |
